# Synaptophysin Chaperones the Assembly of 12 SNAREpins under each Ready-Release Vesicle

**DOI:** 10.1101/2023.07.05.547834

**Authors:** Manindra Bera, Abhijith Radhakrishnan, Jeff Coleman, Ramalingam Venkat Kalyana Sundaram, Sathish Ramakrishnan, Frederic Pincet, James E. Rothman

## Abstract

The synaptic vesicle protein Synaptophysin has long been known to form a complex with the v-SNARE VAMP, but a more specific molecular function or mechanism of action in exocytosis has been lacking because gene knockouts have minimal effects. Utilizing fully-defined reconstitution and single-molecule measurements, we now report that Synaptophysin functions as a chaperone that determines the number of SNAREpins assembling between a ready-release vesicle and its target membrane bilayer. Specifically, Synaptophysin directs the assembly of 12 ± 1 SNAREpins under each docked vesicle, even in the face of an excess of SNARE proteins. The SNAREpins assemble in successive waves of 6 ± 1 and 5 ± 2 SNAREpins, respectively, tightly linked to oligomerization of and binding to the vesicle Ca^++^ sensor Synaptotagmin. Templating of 12 SNAREpins by Synaptophysin is likely the direct result of its hexamer structure and its binding of VAMP2 dimers, both of which we demonstrate in detergent extracts and lipid bilayers.

**Significance Statement:** Synaptophysin is the most abundant protein and a unique constituent of synaptic vesicles, yet it has no known function, due to minimal genetic phenotypes and the lack of biochemical assays. Here, we directly establish using two independent methods that the synaptic vesicle protein Synaptophysin forms a hexameric complex containing 12 copies of the v-SNARE VAMP2. These v-SNAREs assemble into SNAREpins as ready-release vesicles are formed in a fully-defined cell-free system, and do so in two equal waves organized by oligomerization of the Ca^++^ sensor Synaptotagmin. In the absence of Synaptophysin, two waves are also observed, but the number of SNAREpins in each varies widely. We suggest that a single Synaptophysin hexamer in each vesicle symmetrically organizes 6 pairs of peripheral and central SNAREpins, the latter being directly bound to the Synaptotagmin ring. This gives rise to the symmetrical ring-like arrangement of densities observed by cryo-EM tomography under each synaptic vesicle (1, 2).

## Introduction

Although the total free energy available from folding up (“zippering”) of the SNARE complex is ~70 k_B_T (3), most of this exceptionally large amount of energy is utilized before bilayer fusion in order to irreversibly force two docked membranes into close proximity. This can leave as little as ~5 k_B_T (4) available from terminal zippering to overcome the ~30 k_B_T energy barrier (5, 6) for bilayer fusion. Therefore, to achieve the ultra-fast fusion required by synaptic vesicles, multiple SNAREpins must be simultaneously released for terminal zippering. Because they are mechanically coupled by attachment to locally rigid bilayers, as few as 3-6 SNAREpins can potentially cooperate to trigger fusion pore opening in < 1 msec (7).

The surprising discovery from cryo-electron tomography that each such synaptic vesicle sits atop a 6-fold symmetric ring-like arrangement of protein densities (1, 2) implied there are 6 (or potentially a multiple of 6) SNAREpins beneath each ready-release vesicle. The ring likely contains the Ca^++^ sensor Synaptotagmin, as the isolated protein self-assembles into similar size rings (8) and mutation of Synaptotagmin disorganizes the densities under the synaptic vesicles (9). Munc13, a chaperone needed for SNAREpin assembly at synapses (10–13), capable of self-assembly into hexagons between phospholipid bilayers (14) provides one possible mechanism to template up to 6 SNAREpins at a time (15).

The abundant synaptic vesicle-specific protein Synaptophysin (Syp) could provide a second independent mechanism for templating a multiple of 6 SNAREpins. It has long been known that Syp binds the synaptic vesicle v-SNARE VAMP (16–18). Betz et al. (19) and more recently Stowell and colleagues have reported from averaged negatively stained EM images that purified Syp can exist as a hexamer in cholesterol-containing detergent solution (20) where it binds the synaptic v-SNARE VAMP2 in an approximately 1:2 molar ratio (21). This implies that a Syp hexamer could theoretically sequester up to 12 copies of VAMP, offering a simple mechanism that could assemble 12 SNAREpins per vesicle (22). However, direct evidence for this hypothetical complex has been lacking.

Why is such a basic fact as the number of SNAREpins that are assembled in a ready-release synaptic vesicle still unknown three decades after the SNARE complex was discovered and implicated in neurotransmitter release (23–27)? The explanation is that despite enormous advances in imaging technologies, there are still no suitable methods with sufficient sensitivity and resolution to detect and quantify individual molecular components of the release machinery at synapses *in situ*. For example, a careful study of the SNARE assembly chaperone Munc13 using single molecule-based super-resolution imaging could reliably detect approximately 5-10 copies of Munc13 associated with each ready-release vesicle at the neuromuscular junction, but could not quantify their number more precisely (28).

To gain further insights, it is necessary to utilize reconstituted cell-free systems that reproduce ultra-fast fusion (29, 30), and which are compatible with well-established single-molecule methods that enable straightforward counting of directly labeled proteins in buffers and geometries that favor signal detection over backgrounds. Total reconstitution of ultra-fast release minimally requires the synaptic SNAREs, the Ca^++^-sensor Synaptotagmin, the lipid PIP_2_ (for Synaptotagmin function) and Complexin, and has been achieved both between vesicles in suspension (31, 32) and between single-vesicle and planar suspended bilayers (30, 33–35). In these systems, the plasma membrane SNAREs Syntaxin and SNAP25 are pre-assembled into t-SNAREs thereby bypassing the requirement for the specialized chaperoned Munc13 and its partner Munc18 (36, 37). The ultra-fast synchronous release is also reconstituted in a complete system in which SNARE assembly is chaperone-dependent (12, 38).

We recently described a further advance upon the vesicle-suspended bilayer system in which the bilayer is now held within ~ 5 μm of a coverslip above a buffer whose refractive index is raised with a polymer (Iodixanol) to match that of the glass. This chip geometry enables Total Internal Reflection Fluorescence (TIRF) microscopy, whose sensitivity enables the quantitation of single molecules associated with individual docked vesicles whose fate can be followed after the addition of Ca^++^ (39).

In this report, we utilize this advance to directly test the idea that Syp can determine the number of SNAREpins assembling between a ready-release vesicle and its target membrane bilayer. In order to isolate this potential mechanism from the alternative templating mechanism involving Munc13, we utilized the chaperone-free system involving pre-assembled t-SNAREs. We begin by first using related single molecule methods to directly establish the subunit structure of Synaptophysin and its complex with the v-SNARE VAMP2, both in lipid bilayers and in detergent solution.

## Results

### Syp forms hexameric clusters containing up to 12 copies of VAMP2 in lipid bilayers

To test for the formation of oligomeric complexes of Syp and VAMP2 in the native environment of a phospholipid bilayer, we labeled these proteins with different colored dyes, reconstituted them into a common supported lipid bilayer, and determined whether they bound each other and if so in what numerical combinations. By counting single molecules at a very low surface density (~10 Syp molecules per ~1 µm^2^), we could exclude associations by coincidence despite the limits of optical resolution. We chose supported bilayers because the lateral diffusion of proteins is very slow due to absorption to the glass (40, 41), facilitating cluster analysis.

Therefore, recombinant Halo-tagged Synaptophysin was expressed in HEK293 cells, purified from detergent extracts using its 12-his-tag (Fig. S1C), and labeled with Alexa488 chloroalkane. Recombinant VAMP2 (mutated from wild-type by S28C and C103A to allow for selective labeling with Alexa647-maleimide), was expressed in *E. coli* and purified according to its 6-his tag (29, 30, 34). These proteins were then co-reconstituted (Fig. S1C) in a mole ratio of ~ 4 VAMP2 per Syp into small unilamellar vesicles (SUVs) containing phospholipids with 20 mole% cholesterol, included because cholesterol promotes VAMP-Syp association (21, 42, 43). The SUVs were then applied to a clean glass coverslip with 5 mM MgCl_2_ to form supported bilayers (44, 45) and then imaged by TIRF microscopy sequentially with 488 nm (Syp) and 647 nm (VAMP2) lasers. We quantified cluster sizes from both the intensity values and step photo-bleaching curves.

Syp considered by itself (Fig. 1B, top panel) existed in clusters containing 6 ± 2 copies per cluster. When Syp molecules co-clustering with VAMP2 were considered separately, these Syp clusters (Fig. 1B, middle panel) contained an indistinguishable 6 ± 1 copies of Syp per cluster and now also contained 11 ± 3 copies of VAMP2 (Fig. 1B, bottom panel). Under these conditions ~50% of VAMP2 is co-clustered with Syp, the balance being unbound and free.

**Figure 1.**
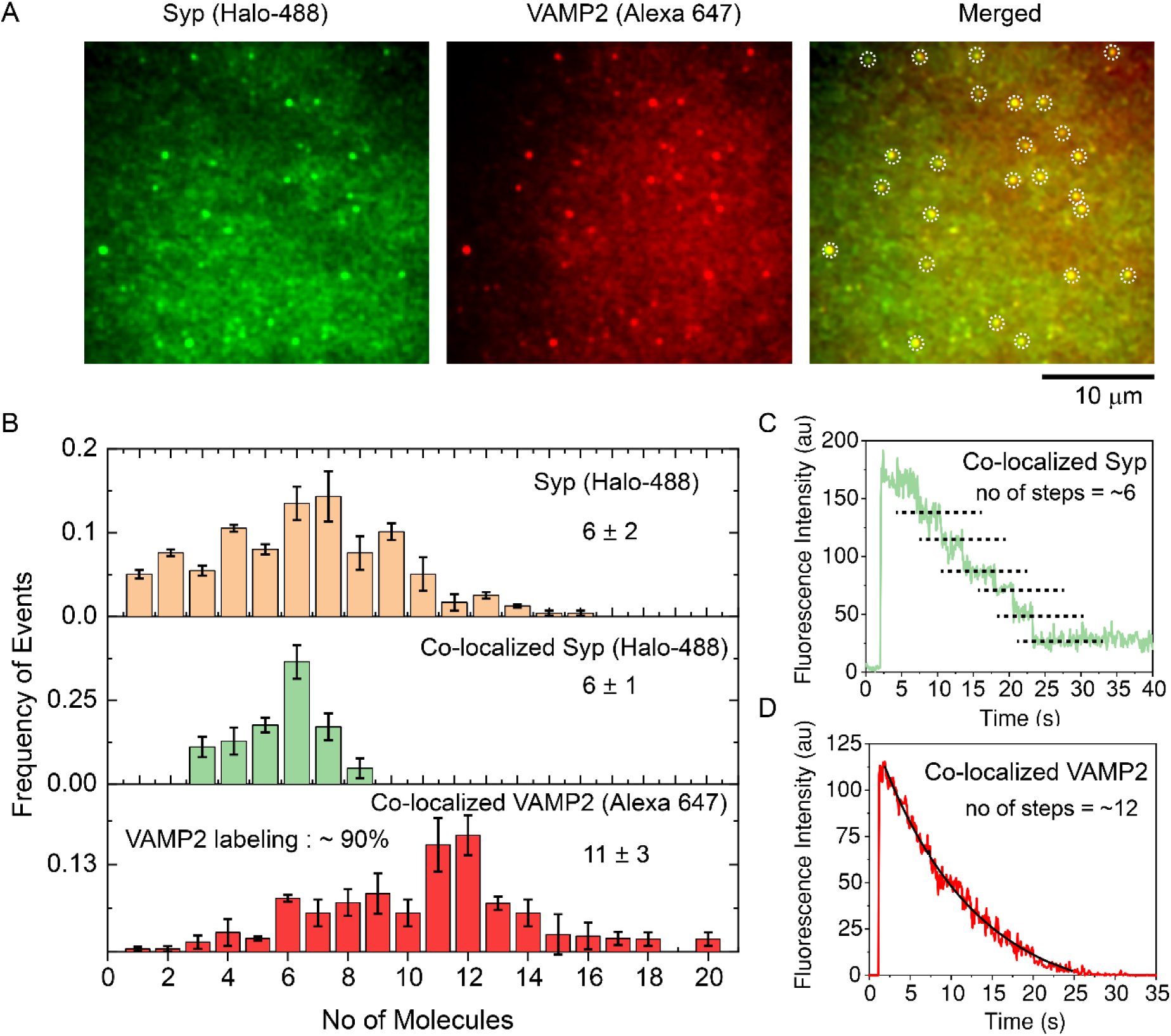
(Syp)_6_-(VAMP2)_12_ complex formation on supported bilayers. Synaptophysin-Halo (Alexa 488) and VAMP2 (Alexa 647) are incorporated into the 20% cholesterol-containing small SUVs to form a supported bilayer onto a glass slide. A) Synaptophysin and VAMP2 imaged with 488 and 633 nm lasers respectively with TIRF microscopy show distinct clusters. Co-localized clusters are marked with a white dashed circle. The scale bar is 10 μm. B) Histograms for Synaptophysin (orange), co-localized with VAMP2 (green) and VAMP2 co-localized with Synaptophysin (red). Mean ± SEM values of all Synaptophysin (6 ± 2), co-localized Synaptophysin (6 ± 1) and co-localized VAMP2 (11 ± 3) cluster size distributions are indicated on the right side of each panel. Black error bars denote the standard deviations. The total no of particles (n) for all Synaptophysin and both co-localized clusters are 1953 and 381, respectively. All the clusters are calculated from three independent experiments. Both Syp-Halo 488 and VAMP2-Alexa 647 labeling percentages were ~95% and ~90%, respectively. C and D) Examples of step photo-bleaching curves for co-localized Synaptophysin (6 distinct steps shown using black dotted lines) and co-localized VAMP2 (total steps ~12) are presented in C and D, respectively. VAMP2 bleaching steps are fitted with an exponential decay equation (black solid line). The fitted initial intensity, i.e. the pre-factor of the fit, is used to derive the total no of steps.

### VAMP2-dependent assembly of Syp into single hexameric particles in solution

From the previous experiment, we can conclude that Syp efficiently forms hexamers in phospholipid bilayers, each of which efficiently sequesters up to ~12 copies of VAMP2, even when these proteins are very sparse. For comparison, the average surface concentrations of VAMP2 and Syp are about 100 and 10 monomers per µm^2^, respectively, as compared to the ~9000 and ~4000 molecules/µm^2^ that can be calculated for native synaptic vesicles from published data (46).

These considerations suggest that complexes between VAMP2 and Syp may also exist in a dilute detergent solution to allow independent single-molecule measurements of stoichiometry following adsorption to cover slips from a dilute solution. Using 1 µM Alexa488-Halo-Syp resulted in clusters containing an average of 3 ± 1 copies (Fig. 2A, top panel). Incubation with unlabeled VAMP2 (4 µM final concentration) produced larger Syp oligomers, now averaging 5 ± 2 copies each (Fig. 2A, middle panel).

**Figure 2.**
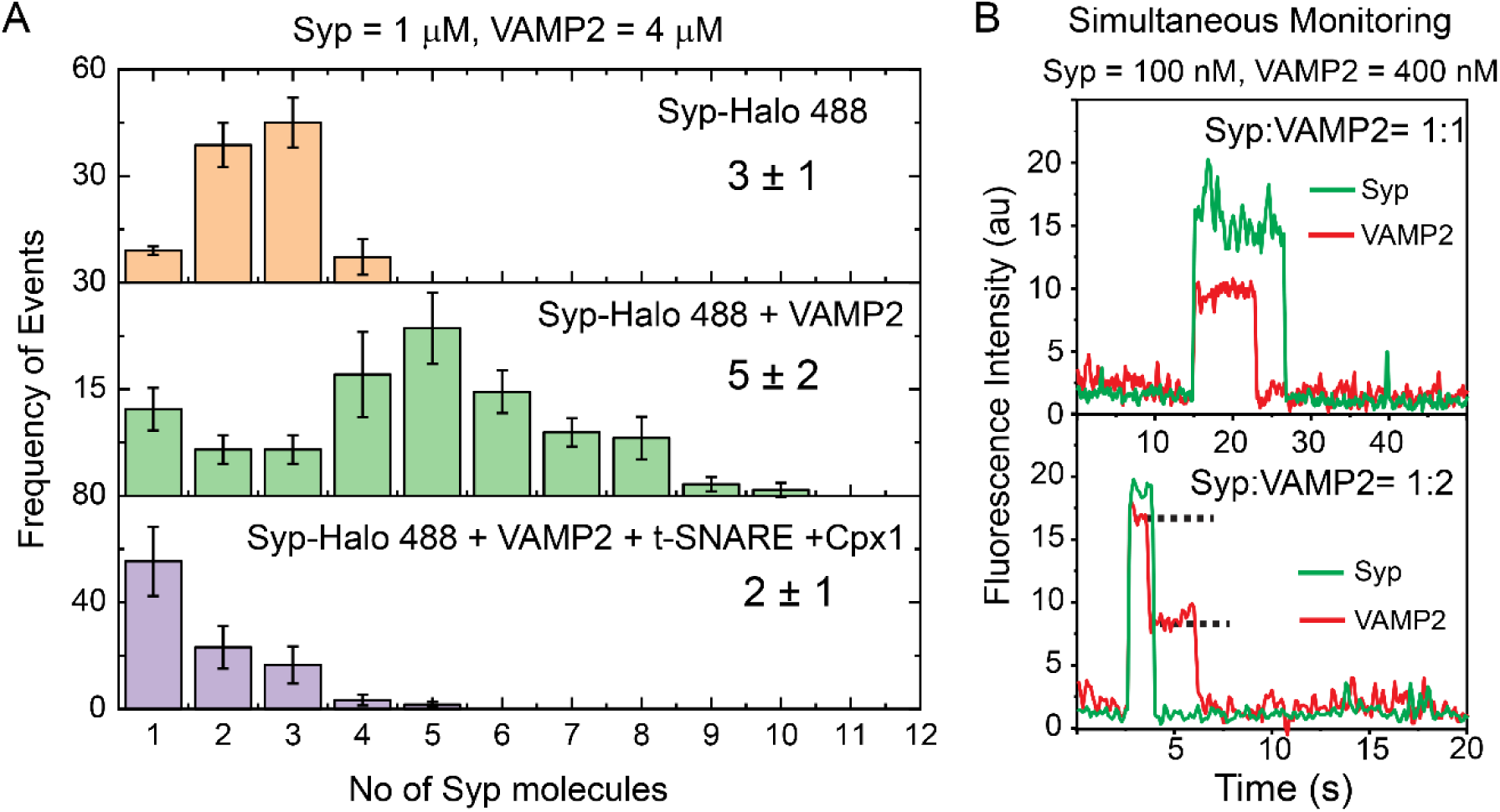
VAMP2-dependent Syp_6_ assembly and the 2:1 complex in solution. A) Distribution of copy numbers of Synaptophysin (Alexa-488) particles when incubated alone (1st panel in orange), after overnight incubation with VAMP2 (1:4 molar ratio, 2nd panel in green) and after a second incubation with t-SNAREs and Complexin (2 μM each) for 30 min at RT (3rd panel in purple). Each protein sample was briefly exposed onto a freshly BSA-coated glass slide, and washed with fresh buffer before imaging with a 488 nm laser. Incubation with VAMP2 shifted the Syp stoichiometry from 3 ± 1 to 5 ± 2. Adding t-SNAREs and Complexin reduced the stoichiometry down to 2 ± 1. n> 721 for each case, N=3 and the error bars represent the standard deviation of the means. B) Time-dependence fluorescence intensity of Syp (Alexa 488, ~95% labeling efficiency) and VAMP2 647 (Alexa 647, 90% labeling efficiency) complexes after O/N incubation at 1:4 molar ratio and deposition onto a glass slide and monitored simultaneously with 488 and 647 lasers. Only particles with the simultaneous landing of Synaptophysin and VAMP2 landed were considered as actual Syp/VAMP2 complexes. Both panels show one example of 1:1 (upper) and 1:2 (lower) Syp: VAMP2 complex. The black dashed line denotes the number of steps for VAMP2. The sudden rise in fluorescence intensity at 15 s (top) and 2.5 s (bottom) corresponds to the landing of the complexes.

This difference suggested that hexamers of Syp not only bind VAMP2 but are also stabilized by VAMP2 binding. If so, sequestering VAMP2 should destabilize the Syp oligomers. This can be achieved by adding soluble t-SNAREs (the 1:1 complex of the cytoplasmic domain of Syntaxin1 and SNAP25) to form SNARE complexes with the VAMP2, and these can be further stabilized by Complexin. This is exactly what happens (Fig. 2A, bottom panel). Now Syp is almost completely dissociated to 2 ± 1 copies, i.e., to monomers.

To independently determine the stoichiometry of VAMP2 binding to Syp in a detergent solution supplemented with 0.1% cholesterol, we used even lower concentrations of Syp-Halo488 (100 nM) to favor dissociation into monomers and also added VAMP2 (400 nM, now labeled with Alexa647). We monitored both molecules coming together at the same spot simultaneously with 100 msec time resolution (Fig. 2B). Under these conditions, we can not only rule out coincidental binding but also determine the stoichiometry of this complex directly with a small number of photo-bleaching steps.

As anticipated, the vast majority of Syp molecules (90 ± 5%) were now only monomers and only 10 ± 5% of these were in a complex with VAMP2. Each such monomer of Syp contained either one (Fig. 2B, top) or two copies of VAMP2 (Fig. 2B, bottom) but never more than 2 copies. The ratio of Syp_1_Vamp_1_: Syp_1_Vamp_2_ in this population of complexes was ~2:1. Given the concentrations of VAMP2 and Syp in this experiment (400 nM and 100 nM) respectively, and assuming a simple two-step independent binding model where [*Syp*] + 2[*VAMP*] → [*Syp*_1_*VAMP*_1_] + [*VAMP*] → [*Syp*_1_*VAMP*_2_], then the unitary binding constant will be in the range of 1-100 µM in the detergent solution.

We conclude that each subunit of Syp can separately bind up to 2 copies of VAMP2 and that a hexamer of Syp in cholesterol-containing lipid bilayers should bind up to 12 copies of VAMP2, in agreement with the measurements using lipid bilayers.

### Single-particle cryo-EM of the native Synaptophysin-VAMP2 complex

Native Syp-VAMP2 complex was purified from rat brains using a previously described protocol (21). The purity of this complex was established with SDS-PAGE (Fig. S3A) and its components were confirmed by western blotting (Fig. S3B). The ratio of VAMP2 to Syp was 1.7 ± 0.2 moles/mole based on the amount of Coomassie staining relative to subunit molecular masses (Fig. S3A). The freshly purified samples were applied to grids and were frozen before being imaged by 200-kV Glacios (Fig. S3C). A total of 731 movies were collected during a single session and the images were analyzed using RELION 3.1 (47). A template-free auto-picking procedure was used for auto-picking, which yielded a dataset of 635,143 particles. Several rounds of reference-free 2D classifications were carried out to remove bad particles. Individual 2D classes of the final set of homogenous particles frequently revealed molecules with clear evidence of 6 subunits (Fig. S3D) as implied by the biochemical analysis (Figs. 1-2) and as previously suggested by negative-stained images (19, 20). We believe that these 2D classes represent a Syp hexamer containing unit that can template 12 SNAREpins. Further processing of these selected set of particles (~ 70,000 particles) yielded 3D density maps with very low resolution which we decided not to pursue. The poor quality of the density maps are possibly due to a combination of reasons such as the presence of a strong detergent micelle of Triton X-100 covering the protein complex, the difficulty in aligning the particles due to the absence of strong cytoplasmic densities for alignment, preferential orientation (lack of side views) as wells as the innate conformational heterogeneity of the purified Syp-VAMP2 complex in solution. Extensive studies of the protein complex reconstituted in non-detergent platforms are required for elucidating a high-resolution structure and revealing the interaction between Syp and VAMP2.

### Counting SNAREpins on docked vesicles

To measure the number of SNAREpins associated with each docked vesicle, we take advantage of the fact that Complexin binds tightly with very low nM affinity to the SNARE complex in 1:1 stoichiometry via its central helix (48–51). In particular, the central helix binds to partially zippered SNARE complexes containing only VAMP2 residues 29-60 (52–54). Therefore, Complexin is expected to bind to half-zippered SNAREpins (52). As with ready-release vesicles *in vivo*, cell-free reconstitutions are clamped beyond this residue (55, 56), thus we can count clamped SNAREpins simply by counting the number of bound Complexin molecules.

To fluorescently label Complexin1 (Fig. S2A and B), we utilize its sole cysteine (residue number 105) which is near the C-terminus domain and is uninvolved in binding to VAMP2 and Syntaxin in the SNARE complex (48, 54). This Cys is labeled with maleimide-conjugated Alexa647, a measured efficiency ranging from ~50-85% (moles/mole) over numerous preparations (Methods).

When the labeling efficiency is high (~85%) and there are small numbers (<7) of SNAREpins on a vesicle, discrete steps of quantal photobleaching are reliably observed and counted (see Fig. 3B for examples). On the other hand, when larger numbers of (>6) SNAREpins are present, discrete step-wise photo-bleaching is rarely observed (Fig. 3C). In this case the number of fluorescent dye molecules can be measured indirectly and less precisely from the prefactor of the exponential decay curve fitting, and considering average unit fluorophore intensity as measured from single bleaching steps observed in vesicles with low copy numbers in the vicinity (Fig. 3C, see Methods for details).

**Figure 3.**
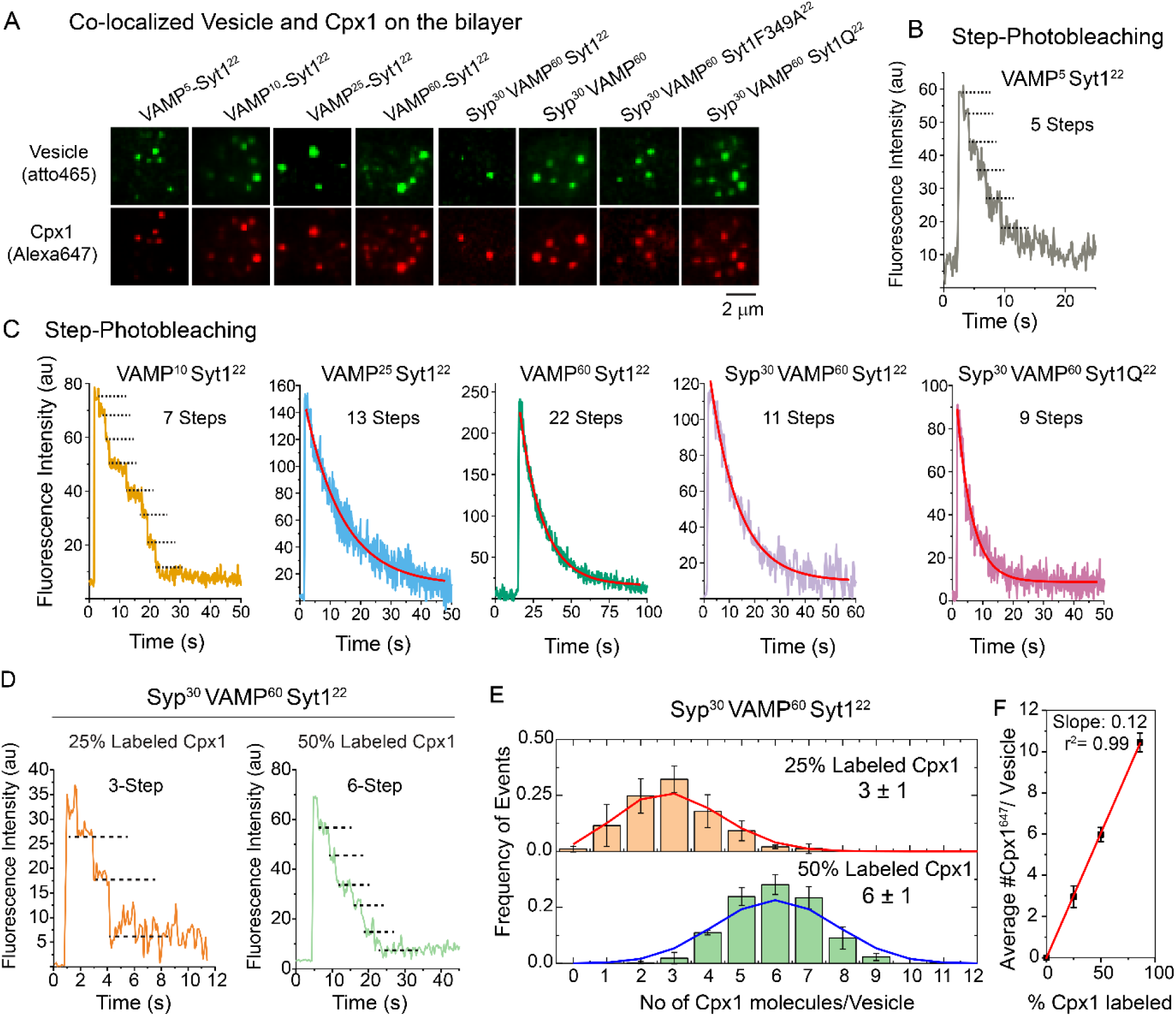
SNAREpin counting under a docked vesicle. A) Representative images for the co-localized docked vesicle and Cpx1 signal on the bilayer. Vesicle and Complexin were respectively labeled with Atto 465 DOPE lipid and Alexa 647-maleimide (85% labeling efficiency). Vesicles and Cpx1 were incubated for 5 min, washed three times and imaged with 488 nm laser and 633 nm laser with 100 ms acquisition time. Scale bar, 2 μm. B and C, Representative traces for Cpx1 photobleaching curves. Cpx1 signals under VAMP2^5^Syt1^22^ (Grey) and VAMP2^10^Syt1^22^ (Yellow) vesicles displayed 5 and ~7 distinguishable photo-bleaching steps, respectively. However, the Cpx1 signal under VAMP2^25^Syt1^22^, VAMP2^60^Syt1^22^, VAMP2^60^Syp^30^Syt1^22^ and VAMP2^60^Syp^30^Syt1Q^22^ vesicles showed smoothly decaying photobleaching curves, indicating higher numbers of photobleaching steps. All the smooth curves were fitted with exponential decay curve fitting to obtain the number of labeled Cpx1 (see method section). D) Examples of step bleaching curves of VAMP2^60^Syp^30^Syt1^22^ vesicles, clamped in the presence of 2 µM Cpx1 with 25% (left) and 50% (right) labeling efficiency. ~3 and ~6 photobleaching steps were observed with 25% and 50% labeled Cpx1 respectively. E) Distributions of the labeled Cpx1 per vesicle. All the 25% (top) and 50% (bottom) labeled Cpx1 numbers were binned from 0 to 12 with bin size 1 and plotted as histograms. Red and blue curves represent the predicted binomial distribution keeping dye labeling at 25% and 50%, respectively and the final SNAREpin number kept at 12. Mean ± SEM values at corresponding labeling efficiency are shown inside the graph. F) The average numbers of labeled Cpx1 per vesicle are plotted with Cpx1 labeling percentages and fitted linearly keeping the intercept at 0. Error bars are standard deviations. For a description of the 85% labeled Cpx1, see Fig. 5B. Slope and linear regression (r^2^) were calculated to be 0.12 and 0.99 respectively. For each case, n>150, N>3.

To add precision when measuring larger numbers of SNAREpins, we utilized defined mixtures of extensively labeled (~85%) and label-free Complexins produced to achieve partial (25% or 50%) labeling. To illustrate why this is useful, consider a case where there are 12 SNAREpins. This vesicle will bind 12 Complexins. If only 25% bear the dye, then on average every such vesicle will have 3 dye molecules, a small number that can be reliably counted by photo-bleaching (Fig. 4D, left panel). However, the actual number will differ from vesicle to vesicle. Some individual vesicles will have bound (at random) 4 dyes and 8 dye-free Complexins; others 2 dyes and 10 dye-free Complexins, and so on. The distribution of labeled complexin follows a binomial distribution with a probability equal to the labeled fraction. Hence, the raw discrete step data for 25%, 50%, or undiluted samples (Fig. 3D and E) can be fitted by a binomial distribution to calculate the average number of SNAREpins for the entire vesicle population more accurately than would have been possible from total intensity analysis. The variation of the mean for the fraction of labeled Complexin follows a straight line with an intercept at 0 and a slope corresponding to 12 ± 1 SNAREpin per vesicle (Fig. 3F).

**Figure 4.**
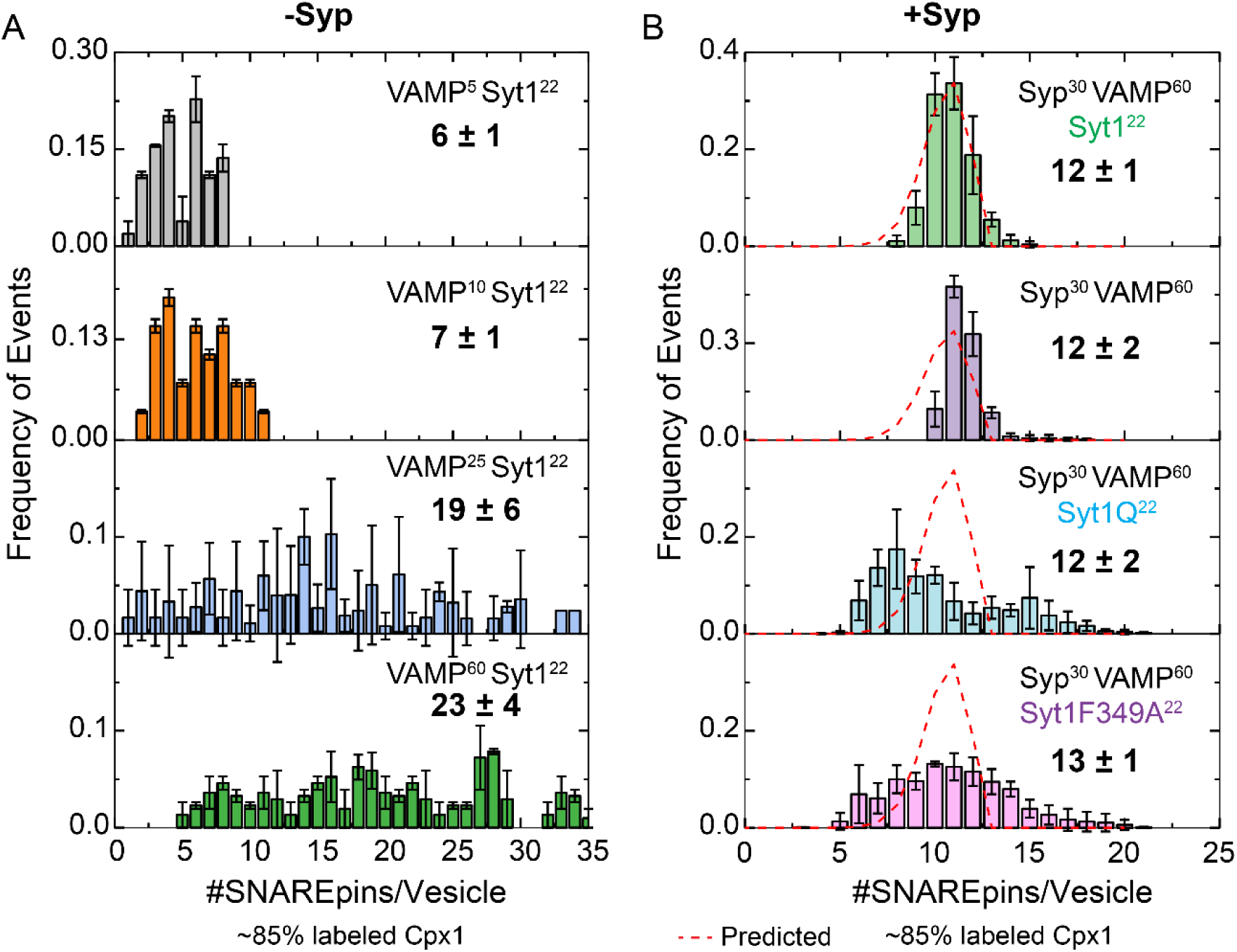
Synaptophysin templates 12 SNAREpins. A) Distribution of the copy numbers of Cpx1 colocalized with docked vesicles without Syp. The total number of Cpx1 molecules under each docked vesicle was calculated from photobleaching steps or total intensity distributions and plotted as frequency histograms with one bin size. Without Syp, an increasing copy number of VAMP2 in the vesicle produces an increasing number of Complexin signals (i.e., SNAREpins). Recombinantly purified VAMP2 and Syt1 were used to reconstitute the vesicles. n>250, N>3 for each case. B) Distribution of the copy number of Cpx1 colocalized with docked vesicles with Syp. With Syp, every vesicle tends to produce ~12 SNAREpins under each vesicle. For Syp^30^ VAMP^60^ Syt1^22^ (green) and Syp^30^ VAMP^60^ (purple) vesicles (n>150, N=3 for each case), natively purified Syp-VAMP complex and recombinantly purified Syt1 (first case) were used. For, the other two cases (Syp^30^ VAMP^60^ Syt1Q^22^ and Syp^30^ VAMP^60^ Syt1F349A^22^), Syp purified from HEK293 cell and recombinant VAMP2 and Syt1were used. The red curve denotes the predicted binomial distribution of the final 12 SNAREpins while the dye labeling percentage was matched with our experimental labeling efficiency (85%). All values in bold letters denote the SNAREpin numbers (average ± SEM) for the corresponding histogram. n> 350, N>3 for each case.

We performed several controls to further establish the specificity of the Complexin signal for counting SNAREpins. As expected, the Complexin fluorescence signal is only detected on the suspended bilayer portion of the chip where it is associated with VAMP2-containing SUVs docked to t-SNARE-containing suspended bilayers (Fig. S2C) and is not observed when vesicles lack the v-SNARE and are instead docked by Synaptotagmin1 (Fig. S2D). This also indirectly rules out the possibility that the labeled Complexin binds to docked vesicles via its curvature-sensing C-terminal amphipathic helix (57, 58) to a detectable extent when diluted for single-molecule studies under our conditions. This was directly established by deleting the amphipathic helix (Complexin1 residues 26-83; Fig. S5). Using this probe and the linear fitting of the mean values (Fig. S5D) we measured 11 ± 2 SNAREpins per vesicle, indistinguishable from the 12 ± 1 measured under the same conditions using the standard full-length Complexin probe (Fig. 4B, top panel). These controls, together with the nature of the results obtained, validate the sensitivity and specificity of the methods we employ to count SNAREpins.

### Synaptophysin limits the number of SNAREpins to twelve

Based on the robust observation of a VAMP_12_-Syp_6_ complex in lipid bilayers (Fig. 1) and the frequent observation of discrete VAMP_1-2_-Syp_1_ subcomplexes in dilute solution (Fig. 2), we hypothesize that the 12 copies of VAMP2 bound to Syp in vesicles will be kinetically advantaged in forming SNAREpins with t-SNARE-containing bilayers. Such a structure, involving 12 SNAREpins, a Syp hexamer, and potentially numerous copies of Syt, would occupy most or all of the interface between the vesicle and bilayer, excluding other SNAREpins from forming. Therefore, our hypothesis predicts that in the absence of Syp, there will be a variable number of SNAREpins, but adding Syp will standardize the number to (theoretically) 12.

We first measured the number of SNAREpins in vesicles lacking Syp (Fig. 5A) but containing variable (5–60) average numbers of externally-oriented VAMP2 and ~22 (on average) copies of externally-oriented Syt and labeled Complexin mixtures. In all of these compositions, vesicles are known from previous work to be efficiently clamped and synchronously released (<13 msec) when Ca^++^ is added (29, 34). With low (5 or 10) copies of external VAMP2, the ready-release vesicles had 6 ± 1 and 7 ± 1 SNAREpins on average, respectively (Fig. 4A, top two panels). With high (25 or 60) copies of external VAMP2, the ready-release vesicles had now had 19 ± 6 (Mean ± SEM) and 23 ± 4 SNAREpins on average, respectively (Fig. 4A, bottom two panels), but the numbers in individual cases are highly variable and can even exceed ~30 SNAREpins per vesicle. These cases seem mainly to involve larger vesicles in the population (Fig. S7) and likely correspond to cases where there are extensive contact surfaces (as distinct from a discrete point of contact) between docked vesicles observed by Cryo-EM (1, 2). For comparison, native synaptic vesicles contain about 70 copies of VAMP2, ~30 copies of Syp (46), and ~15-20 copies of Syt1 (46, 59).

**Figure 5.**
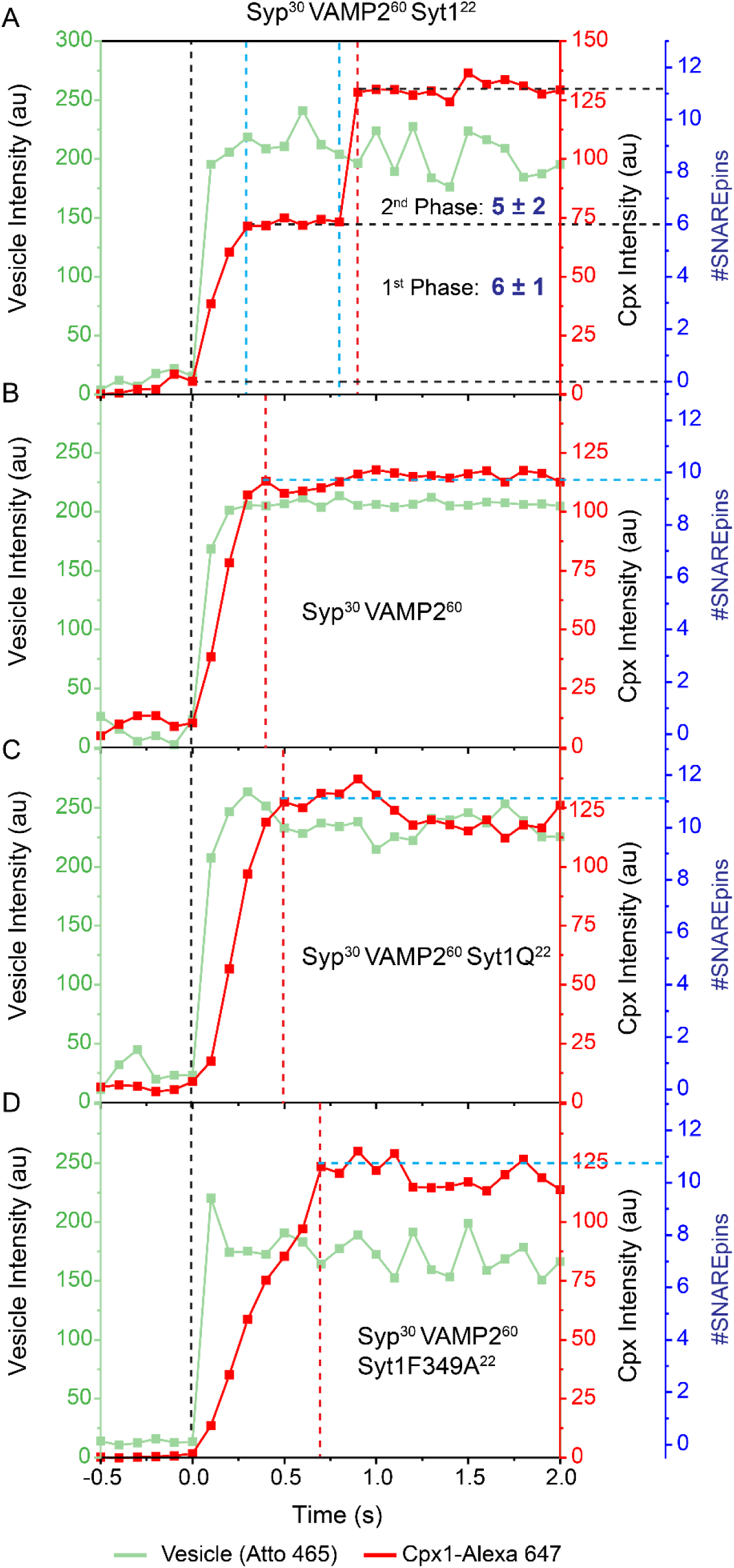
Rates of SNAREpin assembly. A) A representative trace for Syp^30^VAMP2^60^Syt1^22^ vesicles (light green) and Cpx1 (red) assembly are shown. Vesicles and Cpx1 were monitored simultaneously with 488 nm and 633 nm lasers with 100 ms acquisition time. When vesicles dock on the membrane, the Cpx1 signal starts increasing. The time point was adjusted to 0 when vesicles were docked on the membrane, represented by a black dotted line. For these vesicles, Cpx1 showed a two-step assembly. Initially, it increases (1^st^ phase, black and blue dotted lines) and completely stops (lag phase, two blue dotted lines) and then increases again (2^nd^ phase, blue and red dotted line). The left green axis, right red axis and blue axis denote vesicle, Cpx1 signal and the number of SNAREpin formations respectively. The number of SNAREpins formation (blue axis) was calculated from the increase in the Cpx1 signal (right red axis). Black horizontal dotted lines represent the start and end of each phase assembly. The blue horizontal dotted line marks the end of the assembly. B) Syp^30^VAMP2^60^ vesicle showed a one-step rapid Cpx1 assembly. Syp^30^VAMP2^60^Syt1Q^22^ (C) and Syp^30^VAMP2^60^Syt1F349A^22^ (D) *vesicles also showed one-step rapid and slow Cpx1 assembly respectively. Black and red horizontal dotted lines in all panels denote the vesicle arrival and end of SNAREpin assembly respectively*.

When the native Syp-VAMP2 complex purified from rat brains (~30 copies of Syp per vesicle and ~60 copies of VAMP2) was added during reconstitution, the results were dramatic. Now only 12 ± 1 SNAREpins were assembled (Fig. 4B, top panel), compared with the exceptionally broad distribution ranging from 5-35 in the absence of Syp (Fig. 4A, bottom panel). Syp limited the number of SNAREpins to 12 ± 2 even in the absence of Syt (Fig. 4B, 2^nd^ from top). We also introduced recombinantly purified Syp (~30 copies per vesicle) from HEK 293 cells and supplemented with recombinant VAMP2 (from *E.coli;* ~60 copies per vesicle). We observed similar results with a slightly broader distribution.

Importantly, when Syt1 was introduced, the result was a very narrow distribution that closely fits the predicted binomial distribution for 12 SNAREpins considering the experimental dye labeling efficiency of 85% (see Fig. S4). When Syt1 was present, but could not bind SNAREpins (due to mutations in its primary binding site; Fig. 4B, 3^rd^ panel, Syt1Q), or could not assemble ring oligomers (due to an F349A mutation; Fig. 4B, bottom panel), SNAREpin templating by Syp was largely unaffected (12 ± 2 and 13 ± 1 SNAREpins per vesicle, respectively). In control experiments, we confirmed that including Syp (30 copies) in the vesicles along with VAMP2 (60 copies) and Syt (22 copies) did not compromise clamping by Complexin (Fig. S6A) or synchronous release (Fig. S6C-D). Interestingly, Complexin-dependent clamping was observed even in the absence of Syt as long as Syp was present (Fig. S6A). However, Ca^++^-dependent release from this (Complexin plus Syp) clamped state did not occur in the absence of the Ca^++^-sensor Syt.

### Biphasic SNAREpin assembly by Synaptophysin

To study the kinetics of assembly of the 12 SNAREpins templated by Syp we continuously monitored the appearance of immobilized docked vesicles (Atto465) and SNAREpin formation (Complexin-647). Vesicles can be docked by Syt binding to PIP2 in the suspended bilayer and/or by the first SNAREpin to form. Therefore, the vesicle always appears before or together with the Complexin fluorescence. Unexpectedly, we observed that the 12 SNAREpins form in two very distinct phases (Fig. 5A). In the first phase 6 ± 1 SNAREpins are rapidly assembled. Then, following a variable lag phase, a second rapid wave of assembly occurs, resulting in an additional 5 ± 2 SNAREpins. In other words, half the SNAREpins assemble right away, and the other half assemble after a prolonged delay, which averages a highly variable 0.5 ± 1 sec (Fig S9D). Unlike the variable lag phase, the SNAREpins assemble at indistinguishable and rapid rates in the first (34 ± 21 SNAREpins/sec) and second phases (30 ± 21 SNAREpins/sec), corresponding to about 30 msec per SNAREpin.

The basis of the two phases of SNAREpin assembly seems to be in Synaptotagmin, and in particular to binding of SNAREpins to the simultaneously assembling ring of Synaptotagmin. Omitting Syt but retaining Syp produces ~12 SNAREpins but they now assemble rapidly in a single phase (Fig. 5B). Replacing Syt with a mutant that does not bind SNAREpins (Syt1Q) has the same effect (Fig. 5C). Abolishing the ring organization but not SNAREpin binding (Syt1F349A) also eliminates the lag phase but now also greatly slows the rate of SNAREpin assembly (Fig. 5D). Finally, the two-phases of SNARE assembly are even observed in the absence of Syp, but now templating to a limit of 12 SNAREpins is lost (Fig S8 and S9). In the absence of Syp, the first phase produces an average of 10 ± 5 SNAREpins and the second phase has an average of 10 ± 6 SNAREpins for a total of 21 ± 9 SNAREpins (Fig. S9D).

## Discussion

The simplest explanation of our observations regarding the biphasic assembly of SNAREpins is that Syp limits the number of SNAREpins to 12 and Syt ring assembly orchestrates the kinetics. The first 6 SNAREpins assemble around the Syp hexamer while binding 6 individual molecules of Syt. This assembly sterically prevents further SNAREpin assembly until additional copies of free Syt join in to assemble an organized ring containing these “central” SNAREpins. Only when this has been accomplished (during what we observe as the lag phase) can a second set of “peripheral” SNAREpins assemble in the space available outside the ring. When ring assembly is impaired, the lag phase is indefinitely extended and slower untemplated SNAREpin assembly continues as if Syp were absent. When the binding of SNAREpins to Syt is prevented, Syt no longer sterically interferes with the assembly of more than 6 SNAREpins, and the lag phase is abolished but templating to 12 SNAREpins by Syp remains.

Because each vertex of the presumably symmetric Syp hexamer presents 2 copies of VAMP2, it is evident that each copy must be bound differently, either directly to Syp or indirectly as a VAMP2 dimer (21). In either case, their unequal geometries would be expected to afford one copy more reactivity towards t-SNAREs than the other. An alpha-fold prediction of what this structure could look like is shown in Fig. S10. Interestingly, one VAMP2 transmembrane domain (TMD) is predicted to interact directly with Syp 1^st^ and 4^th^ TMD (‘inner’ VAMP2) and the second VAMP2 (“outer” VAMP2) interacts only indirectly when its TMD dimerizes with the TMD of the first VAMP2. We suggest that one copy of Syp at each vertex is pre-determined to assemble the central SNAREpin onto Syt and its ring, while the other (slower-reacting) VAMP2 is destined to form a peripheral SNAREpin. It would seem natural to assume the outer VAMP2 forms the peripheral SNAREpin while the inner VAMP2 assembles into the central SNAREpin. However, alpha-fold offers other types of predictions as well.

In essence, each Syp would asymmetrically bind a VAMP2 dimer. Depending on the stability of the VAMP2 dimers and the stability of Syp oligomers, a mixture of Syp_1_ monomers, Syp_1_VAMP_1_ and Syp_1_VAMP_2_ species would predominate in dilute solution at one extreme, whereas homogeneous VAMP_12_Syp_6_ 18-mers would assemble at high concentrations in lipid bilayers, matching our observations. Moreover, VAMP2 is known to dimerize in cholesterol-containing detergent solutions (43, 60) although not in lipid bilayers as an isolated protein (61, 62) and indirect evidence suggests it interacts with the 3^rd^ and 4^th^ TMD helices of Syp (21, 63).

We have separately suggested that each peripheral SNAREpin is further linked by a bridging molecule of Complexin to the central SNAREpin with which it shares a common Syp vertex to add an energy boost (“turbocharging”) to accelerate fusion pore opening and expansion (*Manuscript under revision*). We have previously provided evidence that central and peripheral SNAREpins (produced in the absence of Syp) are thus linked in this trans-clamping arrangement in which the Complexin accessory helix from one SNAREpin binds to the zippering end of a second SNAREpin (29). The hallmark of trans-clamping is the requirement for an intact accessory helix, also required to stabilize vesicles in the Syp-containing vesicles with 12 SNAREpins (Fig. S11).

It is unclear in our experiments whether the peripheral SNAREpins assemble in the first wave followed by the central SNAREpins, or vice-versa. In any case, the order may differ when t-SNAREs are artificially pre-assembled versus when all three synaptic SNAREs are co-assembled by the chaperones Munc13 and Munc18 (12, 64). In particular, the auto-inhibited Munc13 lateral hexamer is expected to occupy the same space as the Syt ring and is expected to be required for central SNAREpin assembly (14), leading to the prediction (*Manuscript under revision*) that peripheral SNAREpins may assemble before central SNAREpins when the Munc chaperones and Syt are both present.

Synaptophysin has long been known to form a complex with VAMP2 in extracts from synaptosomes (16, 18), but a more specific molecular function or mechanism of action of Syp in exocytosis has been lacking. This is because knockouts of Syp genes in mice have minimal effects on exocytosis (65) as distinct from synaptic vesicle recycling by endocytosis (66, 67). Even a quadruple knockout of the Syp1 and Syp2 along with its homologs Synaptogyrin 1 and 3 had minimal effect in neuronal exocytosis (65). Our new results, from a fully defined cell-free system, stand in bold contrast. Here, Syp has the remarkable and well-defined property of templating exactly 12 SNAREpins under each vesicle. This molecular mechanism is the direct result of its hexamer structure (19–21) and the stoichiometry of VAMP2 binding (21), which we have now directly established to occur in phospholipid bilayers.

To rationalize this clear mechanism of action with Syp’s apparent lack of physiological importance, we need to assume there are independent mechanisms that can adequately control the number of SNAREpins. Munc13 (cooperating with Munc18) chaperone SNAREpin assembly, and in particular Munc13 can assemble between bilayers into at least two hexagonal oligomers (14), lateral hexagons (composed of 6 copies) and a hexamer of upright trimers (composed of 18 copies). It is possible that homeostasis can be maintained by Munc13 alone under many circumstances.

## Materials and Methods

### Materials

The following cDNA constructs were used in this study: full-length VAMP2 (VAMP2-His^6^, residues 1-116); full-length VAMP2^4X^ (VAMP2-His^6^, residues 1-116 with L70D, A74R, A81D, L84D mutations), full-length t-SNARE complex (mouse His^6^-SNAP25B, residues 1-206 and rat Syntaxin1A, residues 1-288); Synaptotagmin (rat Synaptotagmin1-His^5^, residues 57-421); and Complexin (mouse His^6^-Complexin 1, residues 1-134). All mutants including Syt1^F349A^ (F349A); Syt1^Q^ (R281A/E295A/Y338W/R398A/R399A); were generated as described previously (24, 29, 34). Mouse complexin 26-83 was modified with a C-terminal Cys using site-directed mutagenesis. Lipids, 1,2-dioleoyl-sn-glycero-3-(phospho-L-serine) (DOPS), 1,2-dioleoyl-glycerol-3-phosphocholine (DOPC), 1,2-dipalmitoyl-sn-glycero-3-phosphoethanolamine-N-(7-nitro-2-1,3-benzoxadiazol-4-yl) (NBD-DOPE), phosphatidylinositol 4, 5-bisphosphate (PIP2) and cholesterol were purchased from Avanti Polar Lipids (Alabaster, AL). ATTO647N-DOPE and ATTO465 DOPE were purchased from ATTO-TEC, GmbH (Siegen, Germany). Alexa maleimide 647 C2 and Alexa Fluor^R^ 488 Halo ligand were purchased from Thermofisher Scientific (Waltham, MA) and Promega Madison, USA), respectively.

### Protein Expression and Purification

v- and t-SNAREs, Cpx, Syt1 recombinant proteins were expressed and purified as described previously (24, 29, 68). The mouse Synaptophysin gene was cloned as a BamHI/NotI fragment into a modified pCMV6-AN-His vector (Origene, USA) that contains an N-terminal 12xHis tag followed by a PreScission cleavage site. The plasmid was used to transfect Expi293F cells (Thermo Fisher, USA) following the manufacturer’s protocol. Briefly, 1 µg of DNA per 1 ml of cell culture was mixed in a 1:2.5 ratio of DNA: ExpiFectamine reagent for 20 min at RT and then added to approximately 2.5×10^6^ cells with >95% viability. Cells were grown at 37°C under 8% CO2 with orbital shaking for 2 days without enhancers. The cells were centrifuged at 200xg for 10 minutes, washed with PBS and flash frozen. Harvested pellets were stored at −80°C until needed. Expi293F cell pellets (from 50 ml of culture) were thawed and resuspended in 8 ml of 0.25 x PBS with protease inhibitors (Roche, Indianapolis, USA). Cells were lysed using a Dounce homogenizer on ice and then centrifuged in a Beckman SW55 rotor at 100,000xg for 45 min. The pellet was resuspended in 5 mM NaH_2_PO_4_ (pH 6.8), 0.25% cholesterol (Avanti; Ovine Catalog # 700000P) and protease inhibitors. Triton X-100 was added to a final volume of 0.5% (v/v) and incubated on ice for 2 hours with occasional inversion. The lysate was centrifuged at 45,000xg for 30 min to pellet the insoluble fraction and solubilized protein was incubated with 0.125 µl of HisPur Ni-NTA resin (Thermo Fisher, USA) at 4°C overnight on a rotating wheel. The next day, the resin was washed multiple times with 20 mM NaH_2_PO_4_ (pH 6.8), 0.5% TritonX-100, 0.05% cholesterol and 40 mM sucrose. Protein was eluted using the same buffer with 500 mM Imidazole (pH 6.8).

### Synaptic Vesicle Preparation

Synaptic vesicles (SVs) were purified from Sprague Dawley rat brains as described previously (69). Briefly, four rat brains were homogenized in an ice-cold buffer solution (4 mM HEPES, pH 7.4, 320 mM sucrose). The homogenate was centrifuged at 1,000g for 10 min at 4 °C. The pellet was removed, and the resulting supernatant was centrifuged at 15,000g for 15 min at 4 °C. The supernatant was collected and stored at 4 °C while the pellet containing synaptosomes was resuspended in the homogenization buffer and was diluted tenfold with ice-cold ddH_2_O and homogenized thoroughly. The lysate was centrifuged at 17,000g for 15 min at 4 °C. The pellet was discarded, and the resulting supernatant was combined with the supernatant from the previous step and centrifuged at 48,000g for 25 min at 4 °C. The supernatant was further centrifuged at 300,000g for 2 h at 4 °C and the pellet containing SVs was collected. The SVs were immediately resuspended in 3 ml of 5 mM sodium phosphate (pH=6.8), supplemented with 1mM PMSF, 1 µg/ml Pepstatin A and snap frozen in liquid nitrogen and stored in −80 °C until further use.

### Purification of Native Synaptophysin (SYP)-VAMP2 Complex

The Syp-VAMP2 complex was purified from SVs as described previously (21). Briefly, SVs purified from four rat brains were solubilized for 45 min at 4 °C (at 5 mg/ml) in 5 mM sodium phosphate (pH=6.8) buffer supplemented with 1 mM PMSF, 1 µg/ml Pepstatin A, 0.2% Triton X-100, 1% Cholesterol and Protease inhibitor cocktail (Roche). The solubilized Triton X-100 extract was centrifuged at 142,000g for 45 min at 4 °C. The supernatant was applied to a dry hydroxyapatite/celite (2:1 w/w) column and was eluted with 20 ml of 5 mM Sodium phosphate (pH=6.8), 0.2% Triton X-100, 0.1% Cholesterol, 1 mM PMSF, 1 µg/ml Pepstatin A and Protease inhibitor cocktail (Roche, USA). The first five fractions of the flow-through were collected as 1 ml fractions and the rest were collected as 2 ml fractions. A Coomassie gel was run to determine the purity of the Syp-VAMP2 complex in the collected fractions (Fig S1A). The Syp-VAMP2-containing fractions were then pooled and concentrated to 2 mg/ml at 4,200g and 4 °C using 4-ml Amicon Ultra 100-kDa molecular weight cutoff centrifugal filters. The purified protein was stored at 4 °C and was immediately used for cryo-EM sample preparation and reconstitution into liposomes. The purified protein was analyzed by western blot using monoclonal Synaptophysin1 (Synaptic Systems, Germany) and monoclonal VAMP2 (Abcam, USA) antibodies to confirm the presence of the Syp-VAMP2 complex (Fig 1A).

### Cryo-EM Grid Preparation

2.5 µl of freshly purified Syp-VAMP2 complex (1.5 mg/ml) was applied to C-flat-1.2/1.3-2C-T copper grids that had been glow discharged for 10 s in Argon/Oxygen atmosphere using a Solarus plasma cleaner (Gatan, Inc.). The grids were blotted for 2 s with a blot force of −1 at 4 °C under 100% humidity and were flash-frozen in liquid ethane using a FEI Vitrobot Mark IV (FEI). The grids were stored in liquid nitrogen until data acquisition.

### Cryo-EM Data Acquisition and Processing

The frozen-hydrated samples were loaded on the 200-kV Glacios (Thermo Fisher Scientific) with a K2 Summit direct electron detector (Gatan, Inc.). The micrographs were collected in super-resolution mode operated using SerialEM (Version 3) software (70). A total of 731 movies were acquired at a nominal magnification of 36,000x with a physical pixel size of 1.143 Å on the specimen level. The movies were dose fractionated into 40 frames of 0.25 s each at a dose rate of 5.8 electrons/Å^2^/s for a total dose of 58 e-/ Å^2^ over a defocus range between −0.9 µm and −2.0 µm. The image processing was done using RELION 3.1 (47). All the 731 micrographs were motion-corrected and dose-weighted using MotionCor2 (71) with a binning factor of 2 and divided into 5 x 5 patches. The contrast transfer function was calculated with CTFFIND-4.1 (72). A template-free auto-picking procedure based on Laplacian-of-Gaussian (LoG) filter was used for auto-picking, which yielded a dataset of 635,143 particles. Several rounds of 2D classifications using a mask of 70 Å diameter were carried out to remove bad particles and obtain a final set of homogenous particles.

### Liposome Preparation

t-SNAREs and VAMP2 + Syt1 were reconstituted into small unilamellar vesicles (SUVs) using rapid detergent (1% Octyl glucoside) dilution and dialysis method as described previously (24, 29, 34, 35). The lipid composition was ~82 (mole)% DOPC, 15% DOPS, 3% PIP2 and 0.1% ATTO465-PE for t-SNARE SUV and 88% DOPC, 10% PS and 2% ATTO647-PE for VAMP2 and Syt1 SUVs. We used several different lipid: protein ratios in order to reconstitute to desired copy numbers (outward facing) of individual proteins. Copy number of outside-facing proteins was determined using our previously well-established method (29, 34, 35). Recombinant Synaptophysin and VAMP2 (± Syt1) were reconstituted into 50 µl of 3 mM lipids (68% (mole) DOPC, 10% DOPS, 20% cholesterol, 2% ATTO465) that had previously been dried down to a film in a glass test tube using nitrogen gas and then stored under a vacuum for 1 hour to remove any residual chloroform. Lipids were resuspended in protein and buffer (25 mM HEPES, pH 7.4, 400 mM KCl, 1 mM DTT and 0.5% TritonX-100) up to a final volume of 100 µl and then passed through a detergent removal spin column (Pierce, catalog #87777) following the manufacturer’s protocol. Volume was increased to 150 µl and dialyzed against 25 mM HEPES, 120 mM KCl, and 1 mM DTT overnight to remove any residual detergent. The next day, 150 µl of 80% Nycodenz (in the same buffer) was mixed with the sample and then loaded into a 0.8 ml tube (Beckman Coultier, catalog #344090), overlaid with 250 µl of 30% Nycodenz, followed by 50 µl of HEPES buffer. Tubes were centrifuged in an SW55 rotor at 48,000 rpm for 4 hours. Proteoliposomes were collected from the gradient near the buffer/30% Nycodenz interface. Native Syp-VAMP2 complex was also reconstituted following the same protocol except no extra VAMP2 was supplied. Protein reconstitution efficiency was routinely checked with SDS-PAGE and Coomassie staining (Fig. S1). Based on the densitometry analysis of Coomassie-stained SDS gels, the copy number of each protein per vesicle was estimated from trypsin digestion and Coomassie blue staining as described previously.

### Stoichiometry of Syp and VAMP2 on Supported Bilayers

To measure the stoichiometry of the Syp-VAMP2 complex on a supported bilayer, 1 mM (total) SUVs (60 (mole)% DOPC, 15% DOPS, 5% DOPE-PEG 2000 and 20% Cholesterol) were dried with N_2_ gas and kept under vacuum for another 2-4 hr. Dried lipid was dissolved with 25 mM HEPES, pH 7.4, 150 mM KCl, 0.2 mM TCEP and 2% Triton X-100. Pre-labeled and purified Syp-Halo (Alexa 488) and VAMP2 (Alexa 647) proteins were added into the mixture with a protein: lipid ratio of 1:20000 and 1:5000 respectively. 100 mg pre-washed Bio-beads SM2 (Bio-Rad, Hercules, CA) were incubated with that mixture for another 30 min at room temperature with gentle shaking. The liposomes were dialyzed overnight with a 6-8 kD cut-off against a detergent-free HEPES buffer. The liposomes were floated up with a discontinuous gradient of Opti prep. Samples were collected from the top carefully avoiding any Opti prep contamination and further dialyzed for another 2 hr. at 4°C. The supported bilayer was created by Mg^2+^ (5 mM) induced bursting of the liposomes in ibidi glass-bottom chambers (ibidi GmbH, Germany). The bilayer was extensively washed with HEPES buffer. The bilayer was imaged with TIRF (Nikon) microscope using 488 and 633 nm lasers. Photobleaching of the protein cluster was analyzed with ImageJ.

### Syp-VAMP2 Counting on a Glass Surface

Recombinantly purified and labeled Syp-halo 488 (1 µM) was mixed with unlabeled VAMP2 at a 1:4 molar ratio overnight at 4°C in a buffer containing 5 mM NaH_2_PO_4_, pH 6.8, 50 mM KCl, 0.2% Triton X-100, 0.2 mM TCEP, 0.1% Cholesterol. A 5 µl sample was spotted very briefly onto a freshly BSA-coated glass bottom dish (MatTek Corporation, Ashland, MA) and then quickly washed with excess buffer. Syp-halo 488 was imaged with TIRF (Nikon) using a 488 nm laser. The same overnight mixture was further supplemented with 2 µM pre-assembled t-SNAREs and Complexin. After 30 min incubation at room temperature, a similar method was applied to count Syp again. For dual monitoring, both Syp-Halo 488 (1 µM) and VAMP2 Alexa 647 (4 µM) labeled proteins were incubated overnight at 4°C in a buffer consisting of 5 mM Na_2_HPO_4_, pH 6.8, 50 mM KCl, 0.2% Triton X-100, 0.2 mM TCEP, 0.1% Cholesterol. The mixture was diluted (10X) with the same buffer and quickly spotted on the glass bottom dish (BSA coated). Both proteins were monitored using 488 and 647 nm lasers with dual viewer 2 equipped in Nikon TIRF.

### Single Vesicle Docking, Clamping and Fusion Assay

All the single-vesicle fusion measurements were carried out with suspended lipid bilayers as previously described (29, 30, 35). Complexin at a final concentration of 2 μM was added before the addition of vesicles when required. Vesicles (100 nM lipids) were added from the top and monitored for 10 minutes using a 641 nm laser. We employed ATTO647-PE fluorescence to track vesicle clamping and fusion events. Fusion (lipid mixing) was attested by monitoring a burst and then a rapid decrease in fluorescence intensity as the lipid diffuses away. The fate of each vesicle was monitored and plotted as a ‘survival curve’ which represents the fraction of vesicles that docked to the planar bilayer and proceed to spontaneous fusion or immobile docked. After the initial 10 min interaction phase, 100 μM CaCl_2_ was added from the top to monitor the effect of Ca^2+^ on the docked vesicles. Again, the vesicle survivals of the immobile docked vesicles were analyzed, and fractions were binned with 147 ms time. Calcium was monitored as with Ca^2+^-sensor Calcium Green conjugated to a lipophilic 24-carbon alkyl chain (Calcium Green C24) introduced in the suspended bilayer to directly monitor the arrival of Ca^2+^. Details of this method were described in our earlier work. The fusion probability was calculated at the end of 2 min incubation. All experiments were performed at 37°C using an inverted laser scanning confocal microscope (Leica-SP5) equipped with a multi-wavelength argon laser including 488, diode lasers (532 and 641 nm), and a long-working distance 40X water immersion objective (NA 1.1).

### Protein Labeling

Purified complexin proteins (Cpx1^1-134^ and Cpx1^26-83-^Cys) and VAMP2 (S28C, C103A) were incubated overnight with Alexa maleimide 647 C2 dye (Thermofisher Scientific Cat No: A20347) with a 1:5 molar ratio in the presence of 5% DMSO. Syp-Halo protein was incubated with 5 times molar excess halo ligand Alexa 488 (Promega, USA) overnight at 4°C. Excess dye was removed twice using the dye removal column (Thermo Scientific, Cat No 22858) and then protein was loaded into Superdex 200 and concentrated using Amicon Ultra-4 (3K cut off, Merck). The labeling percentage was calculated according to the manufacturer’s protocol. Protein concentration was checked with Bradford assay and dye concentration was measured from the absorbance (Alexa 647 at 651 nm and Alexa 488 at 494 nm) and labeling efficiency was calculated with three repeats.

### Complexin Counting and Rate of SNARE Assembly Measurement

All the complexin counting experiments underneath docked vesicles were performed with TIRF (Nikon) on a reconstituted planar suspended bilayer on the silicon surface. The method to create a suspended bilayer compatible with single-molecule imaging was described in detail elsewhere (39). Briefly, t-SNARE-containing giant unilamellar vesicles (~82% DOPC, 15% DOPS, 3% PIP_2_ and 0.1% Atto 465-DOPE) were prepared using the osmotic shock protocol and burst onto freshly plasma-cleaned Si/SiO2 chips containing 5 µm diameter holes in presence of HEPES buffer (25 mM HEPES, 140 mM KCl, 1 mM DTT) supplemented with index matching 45% Optiprep^(TM)^ (Stemcell Technologies) and 5 mM MgCl_2._ After waiting 20 minutes, the suspended bilayer was extensively washed with HEPES buffer containing 1 mM MgCl_2_. Before adding any vesicles, the bilayer was completely bleached and then 2 µM total complexin (with corresponding labeled protein) was added and mixed well. Then 100 nM SUVs (63% DOPC, 15% DOPS, 20% Cholesterol and 2% Atto 465-DOPE) containing different proteins were added and monitored both vesicle and complexin simultaneously with 488 and 633 nm solid-state laser respectively using dual viewer 2. To generate more statistics on the vesicle and complexin counts, excess complexin was removed by washing three times with HEPES buffer. Both vesicle and complexin were imaged and complexin signals were subjected to photobleaching using a 633 nm laser with 100 ms acquisition time. All the data were analyzed with ImageJ. The number of complexin signals underneath a vesicle was calculated from the steps of the photo-bleaching curves. However, we could not precisely determine the number of steps when the number of molecules was higher than 5-7. The photobleaching curves were fitted with an exponential decay curve with Eq 1.

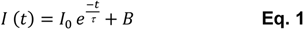

Where I(t) denote the intensity at time t, I_0_ is the initial intensity before bleaching, τ is the time constant, and B is the background. Hence, the copy number, N can be obtained from Eq 2.

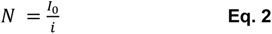

Where i is the average unit intensity of a single fluorophore as determined from the small clusters. For rate of assembly, vesicle and corresponding complexin signal were plotted to monitor the SNAREpin formation. Realtime SNAREpin formation was calculated from the complexin signal using the **Eq 3**.

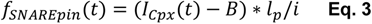

I_Cpx_(t) represents the complexin intensity at time = t, B = background intensity, l_p_ =labeling percentage factor and i, average intensity for one complexin molecule. All the complexin counting data was converted to histograms with 1 bin size. The prediction was calculated using a binomial distribution. We assume there are exactly *N_S_* SNAREpins that form in the contact between the vesicle (Syp^30^ VAMP2^60^ Syt1^22^) and the membrane, and that a single Cpx binds to each SNAREpin. Only a fraction f of the Cpx molecules is labeled. Since Cpx binds randomly to SNAREpins, the number of fluorescent dyes, n, follows a binomial distribution:

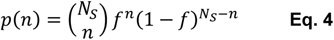

Using N_S_ = 12 and the three labeling fractions used, approximately 0.25, 0.5 and 0.85, the dye copy number distributions can be directly predicted and compared with the experimental ones.

## Acknowledgements

We thank Dr. Kirill Grushin for helping in cryo-EM data collection and processing and generating the structural models. We also thank Dr. Seong Lee for his help in isolating rat brains. This work was supported by the National Institute of Health (NIH) grant DK027044.

## Author Contributions

M.B, A.R, J.C and R V.K.S performed all the experiments; M.B, A.R, S.R and F.P analyzed all the data; M.B, A.R, S.R, F.P and J.E.R designed research; M.B, A.R, J.C, F.P, and J.E.R wrote the paper. All authors read and revised the manuscript.

## SI Appendix

**Figure S1.**
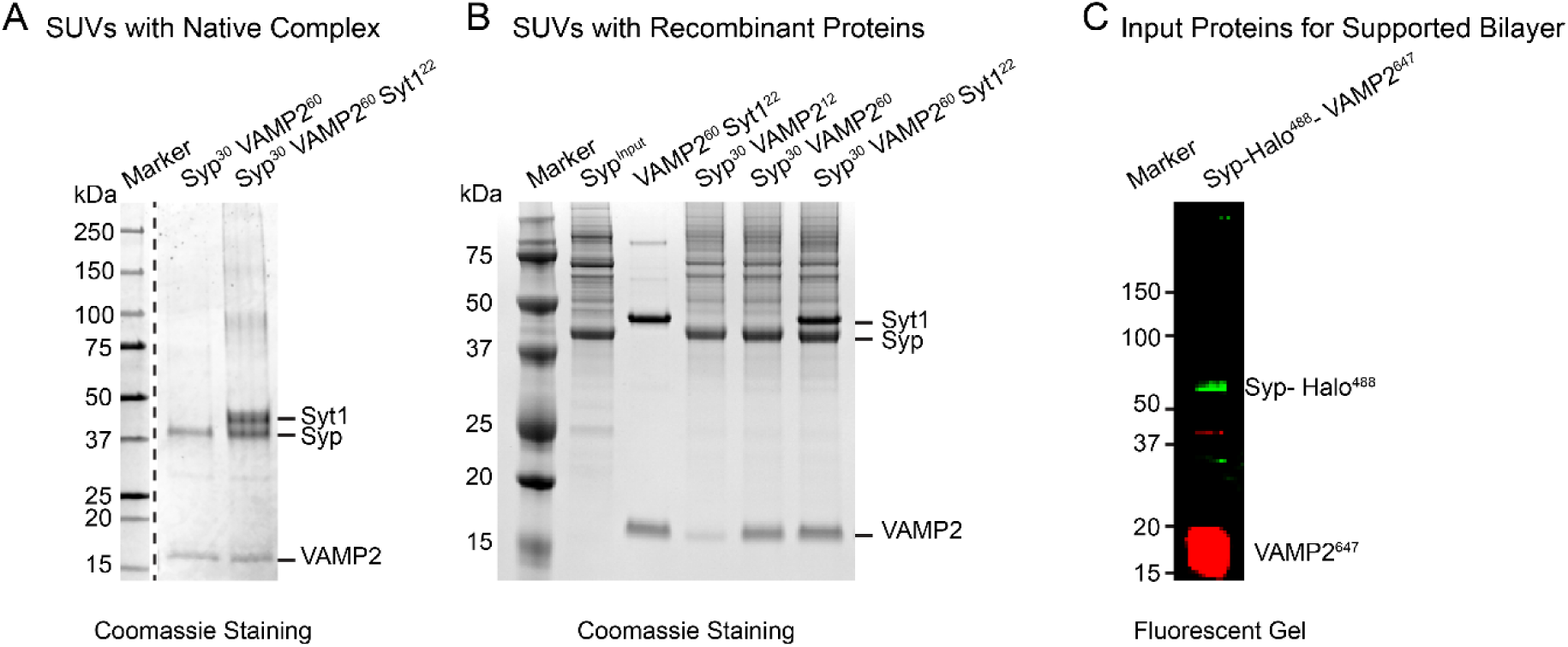
SUV reconstitution with proteins. A) Native Syp-VAMP2 complex was reconstituted into SUVs and floated up through a Nycodenz gradient. No extra VAMP2 was added, however, recombinantly purified Syt1 was loaded into the vesicles if mentioned. B) All recombinantly purified proteins (VAMP2 and Syt1 from E.coli, Syp from Expi293 cells) were reconstituted into the SUVs. SDS-PAGE was run after floating up all the SUVs and staining with Coomassie Blue. Both Syp (native and recombinant) proteins run a little higher than 37 kDa. On the left side, the molecular weights of the markers are noted. Syp input denotes purified Syp protein as input, and superscript numbers represent the copy number (outward facing) of the proteins in each vesicle. C) SDS-PAGE was run with pre-labeled proteins (Syp-Halo with Alexa 488 and VAMP2 (S28C, C103A with Alexa 647 maleimide) which were used to reconstitute SUVs for supported bilayer experiments and fluorescent image of that gel was taken with 488 and 633 nm lasers.

**Figure S2.**
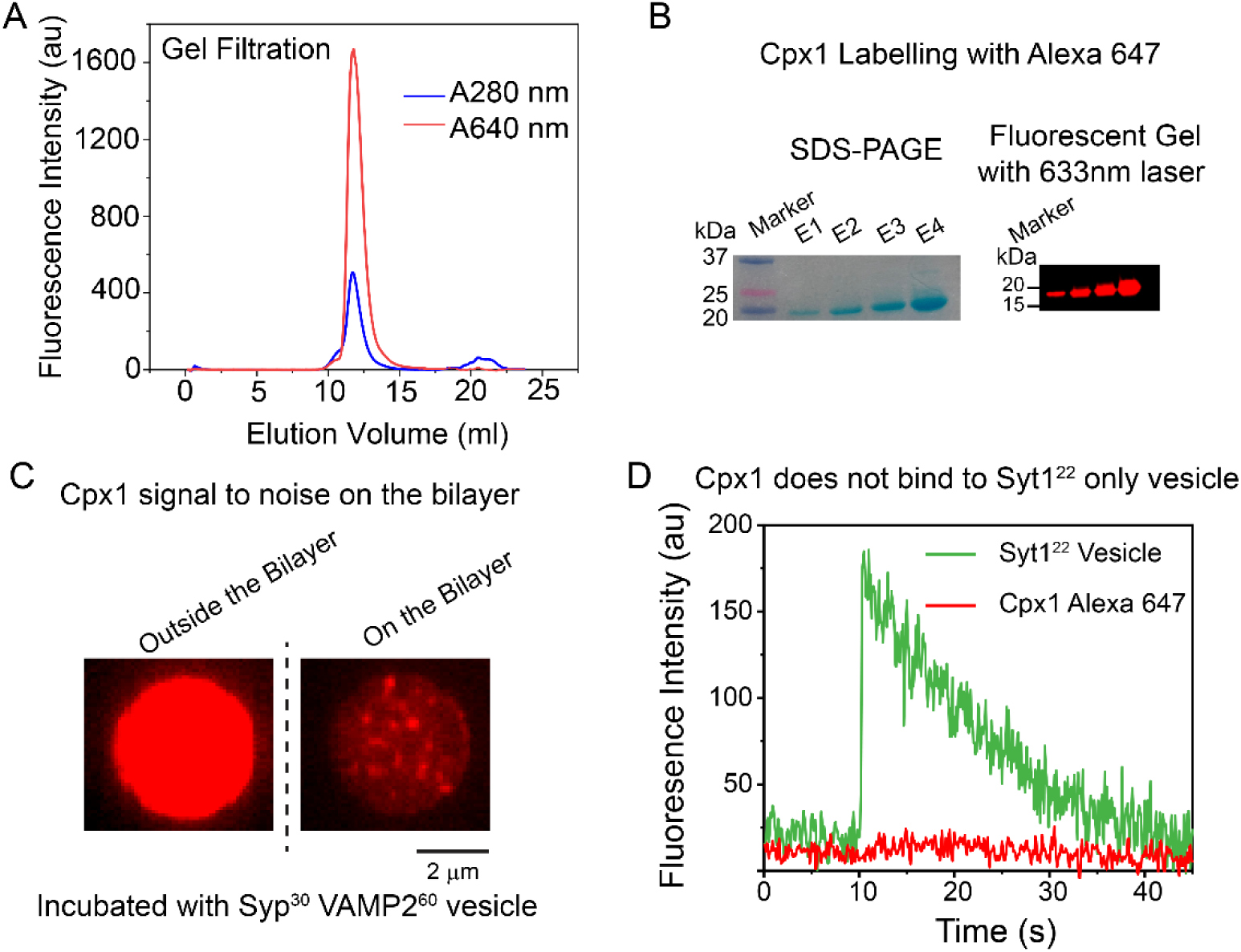
Cpx1 labeling method. A) Complexin 1 was labeled with Alexa 647 maleimide and purified 2X times with a dye removal column and then load onto the Superdex 200 gel filtration column. Both absorbances at 280 and 640 nm were monitored. B) Gel-filtered protein was checked with 4-20% SDS-PAGE. Gel was imaged with natural light (left panel) and fluorescent gel imager with 633 nm laser (right panel). C) ~85% Alexa 647 labeled Complexin has incubated with Syp^30^ VAMP2^60^ vesicle at 2 μM. Complexin adsorbed non-specifically to the silicon surface outside the bilayer (left side) while on the bilayer (right panel), Complexin showed distinct clusters with the high signal-to-noise ratio. Scale bar, 2 μm. D) ~22 copies of Syt1 were reconstituted into a vesicle and incubated with ~85% labeled Complexin. Vesicle (with 488 nm laser) docking and Complexin (with 633 nm laser) arrival were measured simultaneously. Vesicle was docked on the membrane plausibly via the polybasic patch, however, no discernable Complexin signal was observed under that vesicle. After docking, the vesicle was monitored until it slowly bleached.

**Figure S3.**
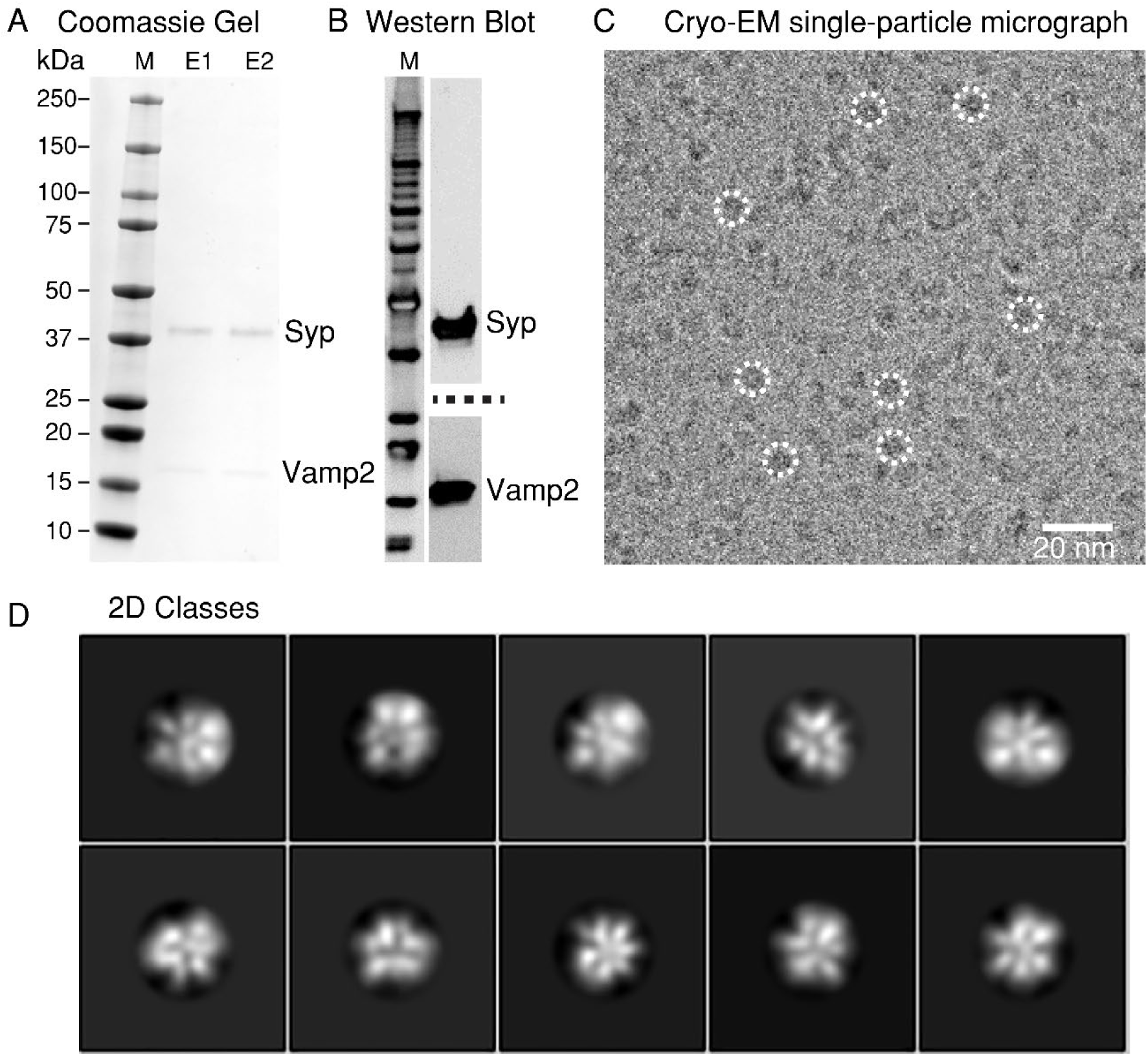
Evidence of hexameric organization of Syp_6_ VAMP_12_ from single-particle cryo-EM. A) SDS-PAGE after purification from rat brain. Two different eluted fractions are shown, and markers are noted on the left side. B) Western blot against rat Syp and VAMP2 antibodies. The left side represents the western blot markers. C) Single-particle Cryo-EM micrograph: White dotted lines denote the complexes picked for further processing, scale bar 20 nm. D) Representative 2D classes of the native Syp-VAMP2 complex showing molecules with six subunits.

**Figure S4.**
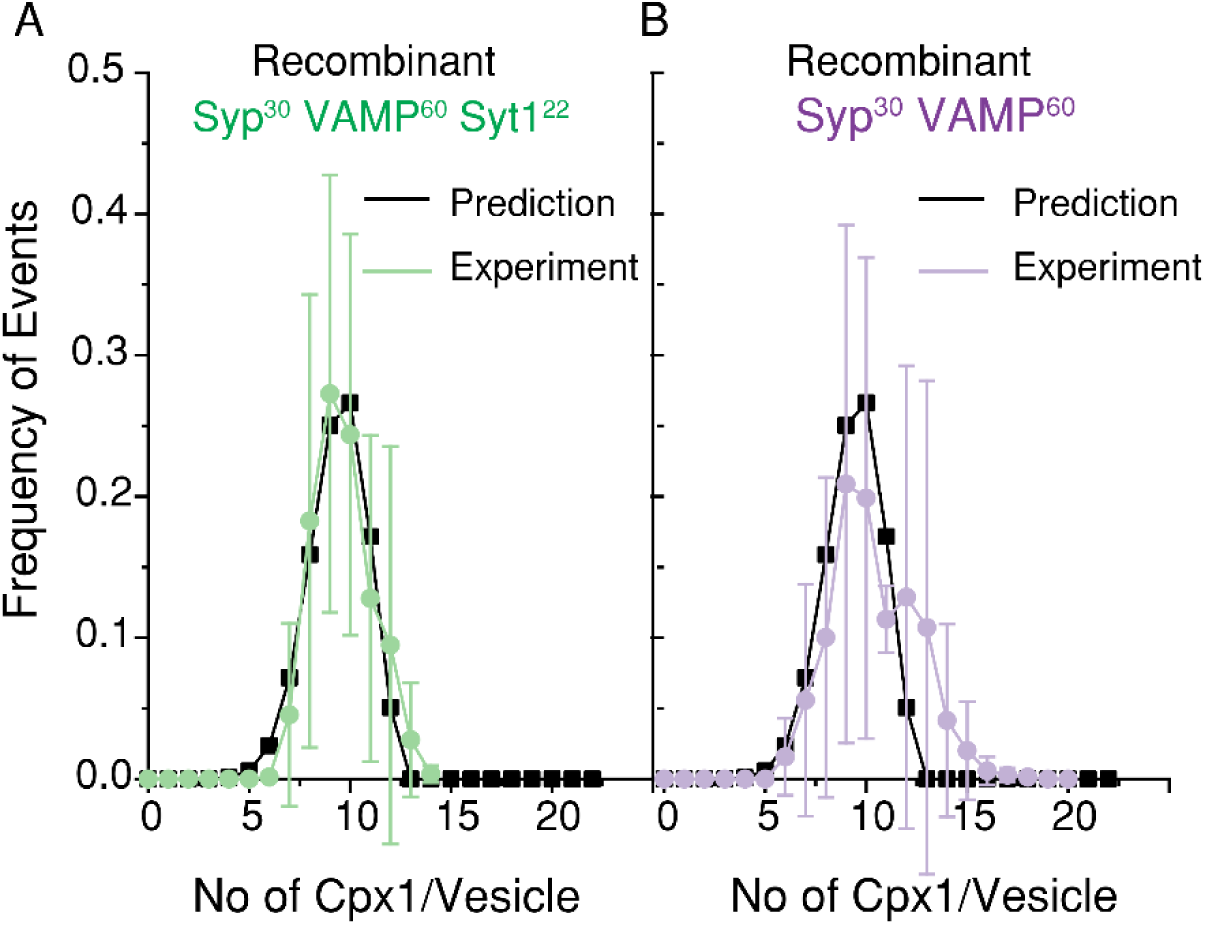
Comparison of SNAREpin formation between recombinantly purified Syp-containing vesicle and prediction. Vesicles containing recombinant Syp and Syt1 together (A, green) produce a narrow distribution (Mean ± SD: 12 ± 1) while in the absence of Syt1 (B, purple), vesicles produce a wider distribution (12 ± 2). The black curve in each case denotes the predicted values considering a binomial distribution with the final 12 SNAREpins with 85% labeling (experimental value) efficiency.

**Figure S5.**
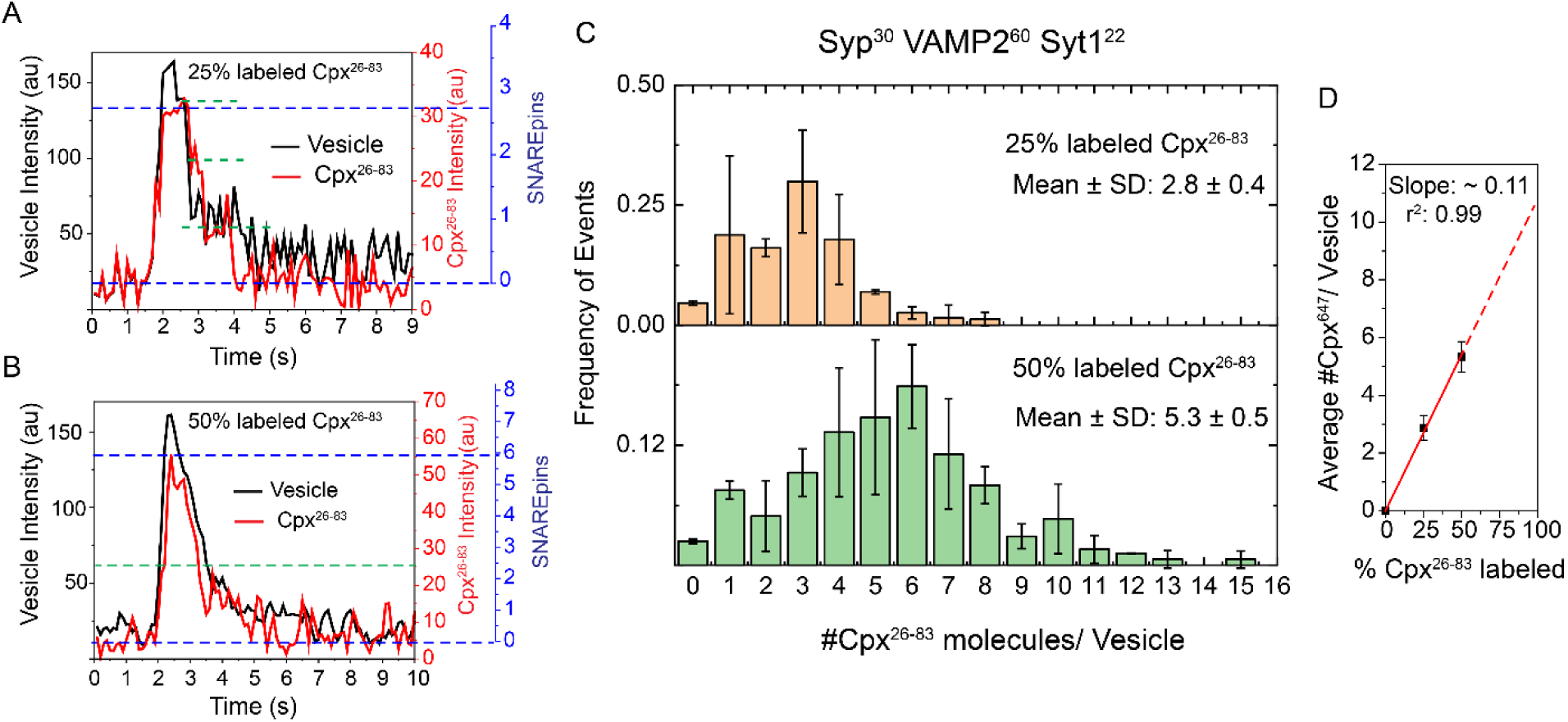
SNAREpin counting with Complexin central helical domain (26–83). Vesicle (labeled with Atto 465 DOPE) clamping, and Complexin^26-83^ (Alexa 647) arrival were simultaneously measured using 50 percent 488 nm and 20 percent 633 nm lasers, respectively, with a 100 ms acquisition rate. A) At 25% doping, vesicle and Complexin arrived simultaneously, however, Complexin showed 3-step photobleaching. The right axis (red) shows the intensity of Complexin, and the blue axis represents the number of SNAREpins. The number of SNAREpins (equivalent to the number of Complexin molecules) was calculated by dividing the Complexin unit intensity value. B) Similarly, vesicle clamping and Complexin were monitored with 50% labeling efficiency. Both vesicle and Complexin showed smooth bleaching. From the Complexin intensity (red, right axis) value, the number of SNAREpins was calculated (represented in right axis, blue). C) From all the step bleaching curves, the number of Complexin molecules under a stably docked vesicle was calculated and binned from 0-16 with 1 bin size. 25% (upper panel in orange) and 50% (lower panel in green) of labeled data were plotted and fitted with Poisson’s distribution. For 25% and 50% label Cpx ^26-83^, mean ± SD were 2.8 ± 0.4 and 5.3 ± 0.5 respectively. n>300, N>3. D) Average Complexin numbers (with SD) were linearly plotted fitted keeping the intercept at 0. From the slope (10.8), the final number of SNAREpin was estimated to be ~11. Linear regression (r^2^) value for the liner fitting was 0.99.

**Figure S6.**
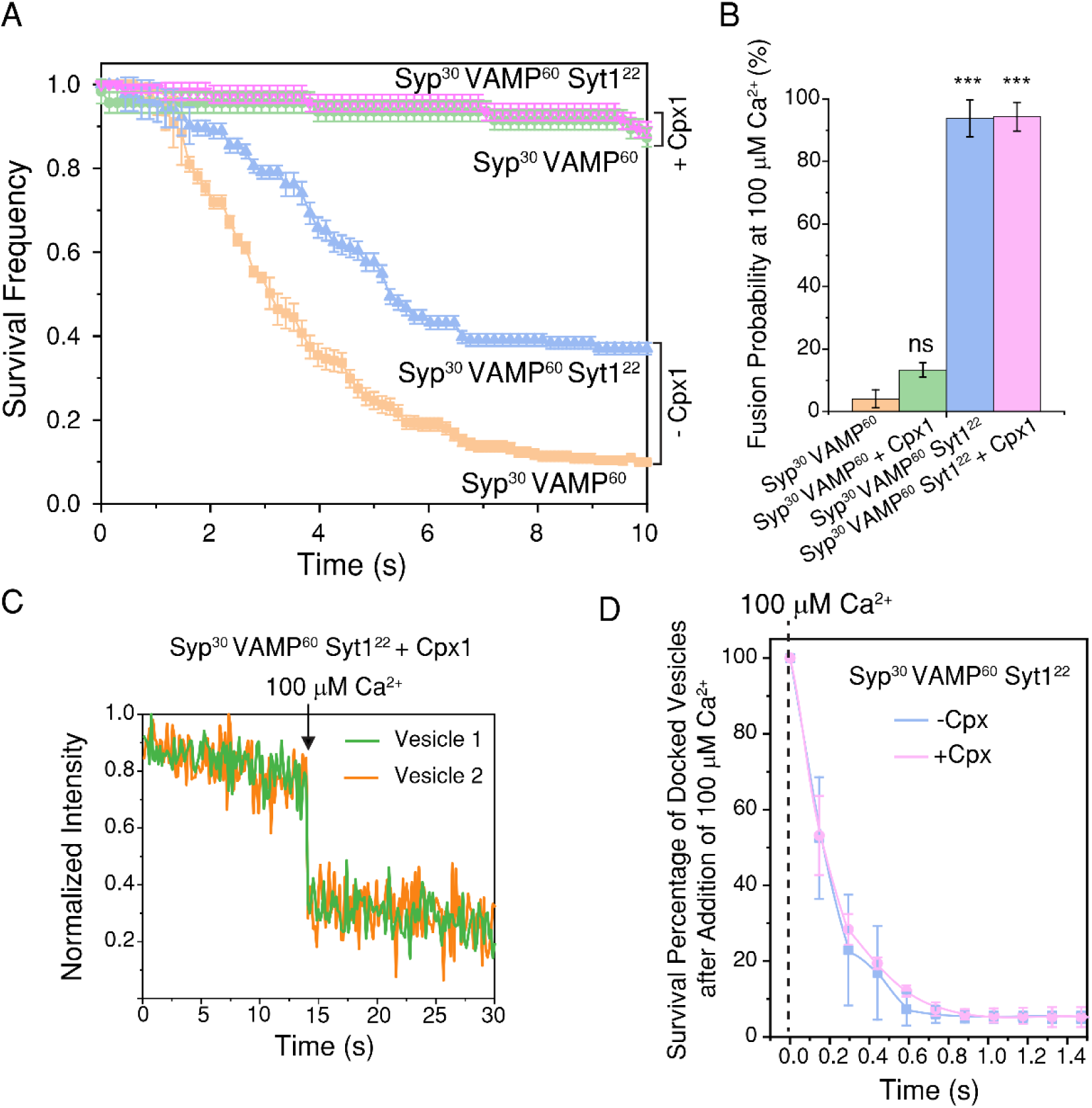
Vesicles containing Syt1 are clamped and ready to release. A) Survival percentages (Kaplan-Meier plot) were calculated at 10 s post docking with different types of vesicles. Vamp^60^-Syp^30^ (orange) vesicles produce a 2-6 sec delay before undergoing spontaneous fusion whereas the addition of 2 μM Cpx1 (green) produces ~95% stably immobile docked vesicles. The addition of ~22 copies of Syt1 into VAMP^60^-Syp^30^ vesicles (blue) generated ~40% stable clamp, however, 2 μM Cpx1 clamped ~95% vesicles (purple). Survivals are calculated when the vesicle appears on the bilayer (t=0) and diffuse freely until either spontaneously fusing or stably docked till photobleached (t=t). All the images are acquired at a 147 ms frame rate. Each curve represents mean ± SD from 3-4 independent experiments (N) and for each case, total Vesicle number (n)> 150. B) Probability of fusions (lipid mixing) is measured 2 min after the addition of 100 μM Ca^2+^. Only Syt1 containing vesicles (Blue and purple) fused with ~95% probability. *** represents P< 0.001 whereas ‘ns’ means not significant. N>3 and n> 100 for each case. C) Representative traces for the synchronous fusion of Vamp^60^-Syp^30^ Syt1^22^ vesicles with Complexin. Two individual vesicles in one hole are undergoing ultra-fast fusion synchronously upon the addition of 100 μM Ca^2+^. In these experiments, vesicles contain 2% Atto647 DOPE to track and upon arrival of Ca^2+^, lipids burst and quickly diffuse away which results in a sharp drop in intensity. D) Lipid mixing was monitored after 100 µM Ca^2+^ addition and survival percentages of the clamped vesicles were calculated after binned with 147 ms (frame rate for these experiments) interval. In both conditions (presence and absence of Cpx), t_1/2_ values for the lipid mixing were calculated to be around ~90 ms.

**Figure S7.**
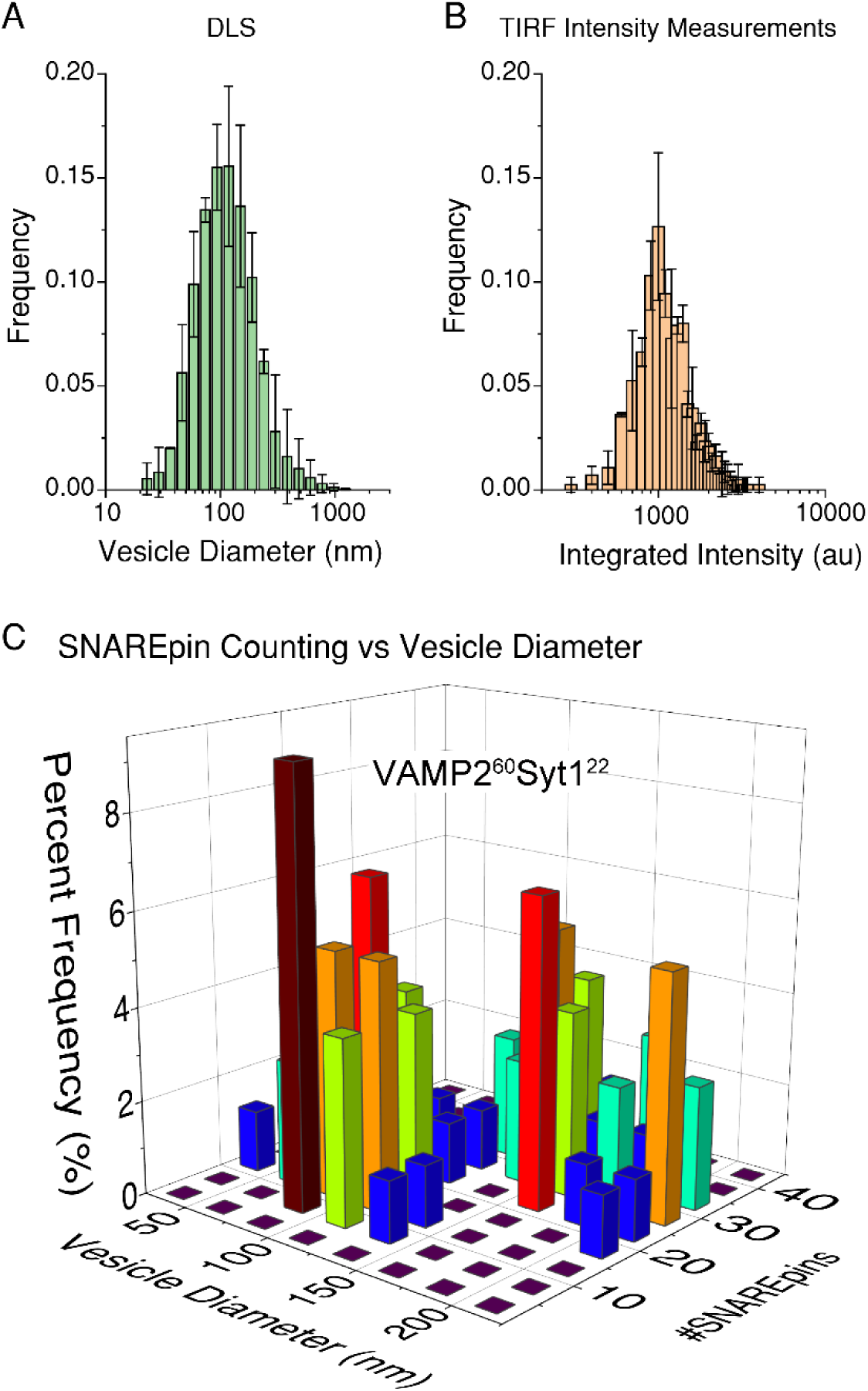
Variation of SNAREpin numbers with vesicle diameter in the absence of Syp. A) Dynamic light scattering (DLS) profile of the VAMP2^60^ Syt1^22^ showing an average diameter of 134 ± 22 nm. N=3 B) The same vesicle raw integrated intensity was measured on the bilayer keeping the ROI fixed at 1.6 μm x 1.6 μm (2.56 x 2.56 pixels). The average integrated intensity was measured to be 1273 ± 100. n> 440, N=3. C) The vesicle intensity was converted into the diameter and plotted against the number of SNARE pins formed underneath that docked vesicle. Vesicle diameter and SNAREpin counts are binned with 20 nm and 5 respectively and plotted with frequency in a 3D plot. n> 75, N=3.

**Figure S8.**
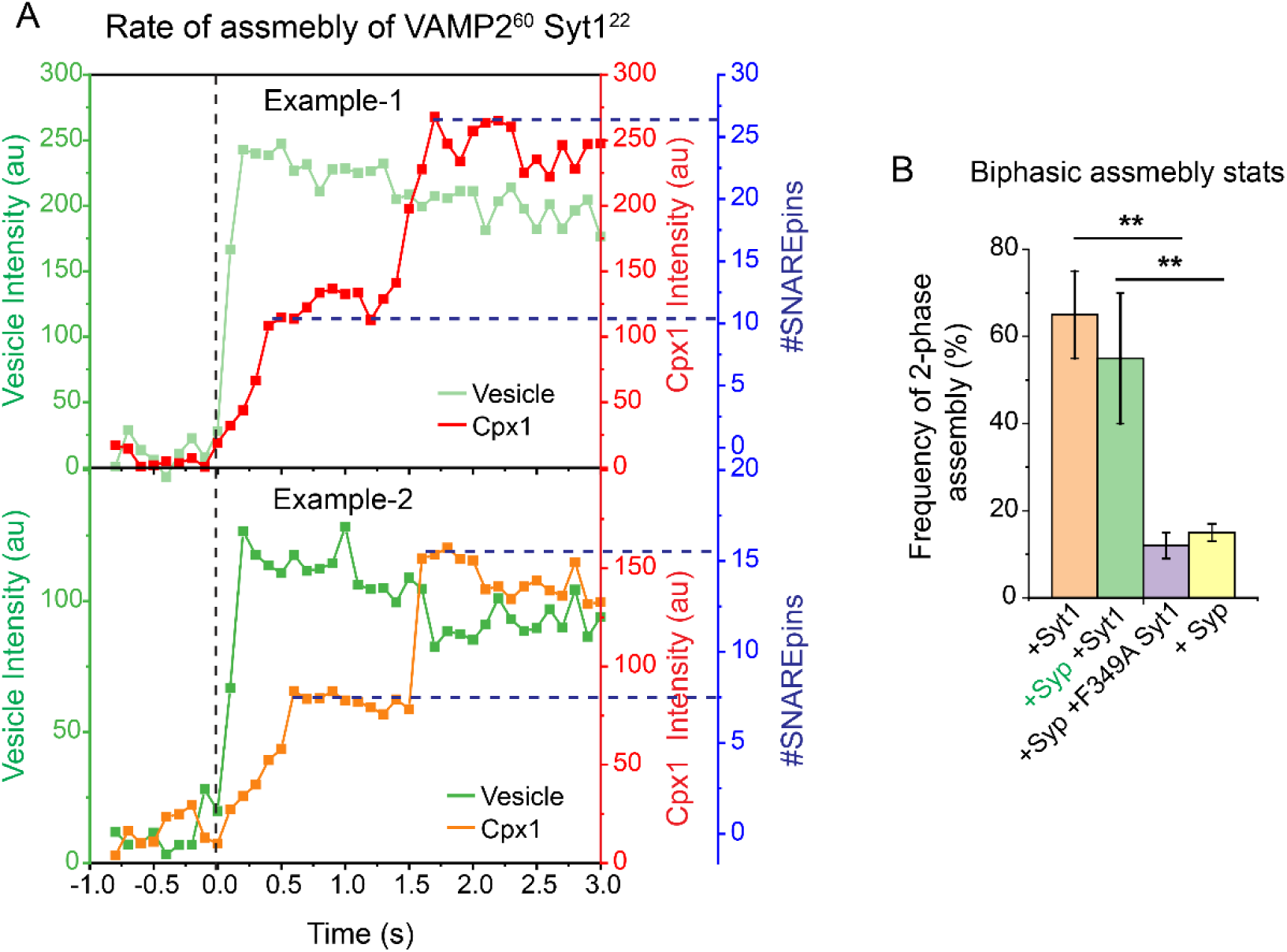
Biphasic assembly in the absence of Syp. A) Two representative traces for VAMP2^60^Syt1^22^ vesicles (light and deep green) and Cpx1 (red and orange) assembly are shown. Vesicles and Cpx1 (85% labeled) were monitored simultaneously with 488 and 633 nm lasers with 100 ms acquisition time. VAMP2^60^Syt1^22^ vesicles showed a two-step Cpx1 assembly. Examples 1 and 2 correspond to one big and small vesicle respectively. The left axis (green), right side red axis and blue axis denote vesicle, Cpx1 signal and the number of SNAREpin formations respectively. Blue and orange horizontal dotted lines represent the end of the 1^st^ phase assemblies for 1^st^ and 2^nd^ examples respectively. B) The percentages of the 2-step assembly processes shown by different vesicles are calculated. All vesicles contained 60 copies (outward facing) of VAMP2 along with the marked proteins. ** p>0.01.

**Figure S9.**
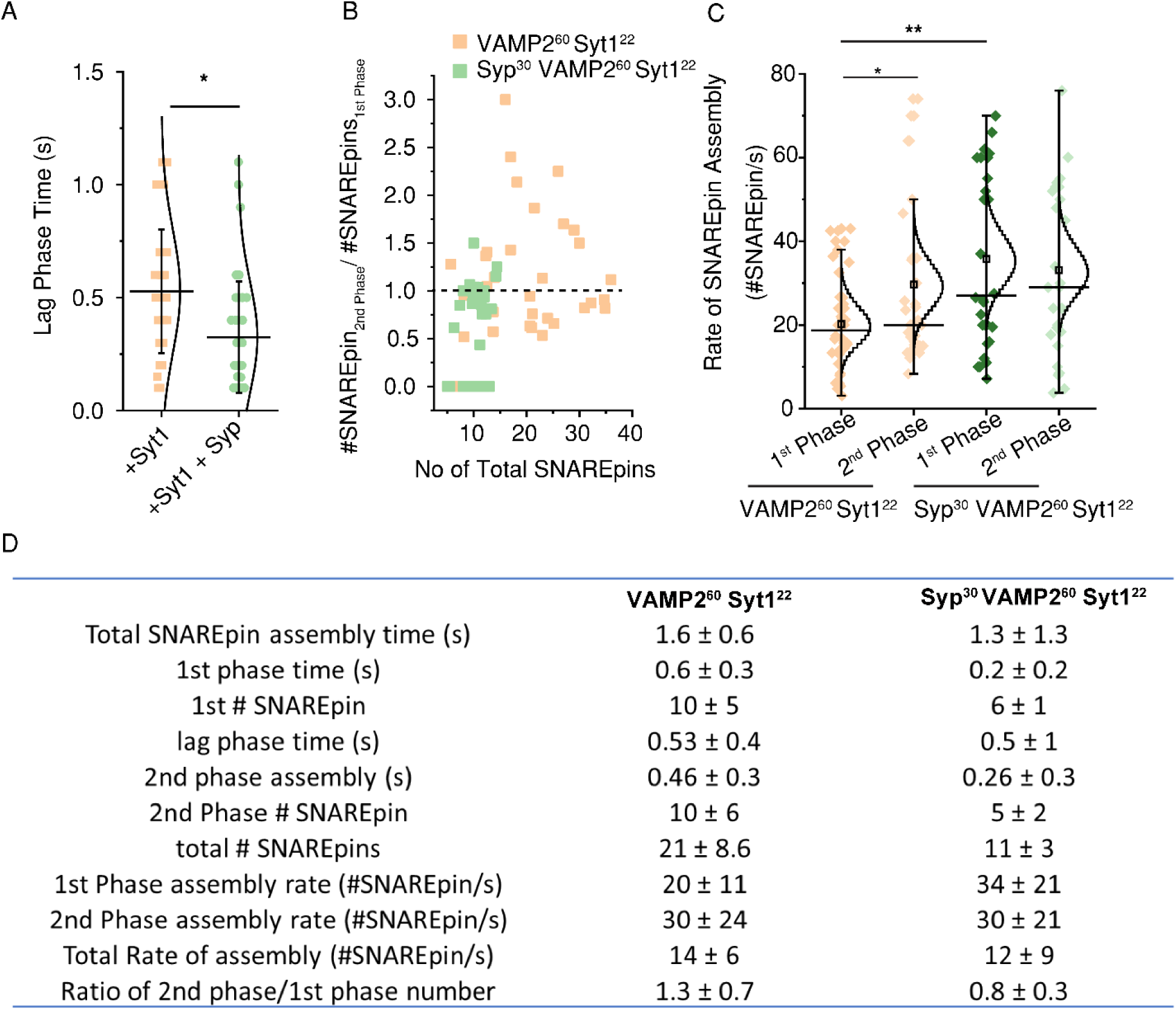
Kinetic analyses: The effect of Syp on SNAREpin assembly. A) Lag phase time taken by VAMP2^60^Syt1^22^Syp^30^ and VAMP2^60^Syt1^22^ vesicles. Each distribution was fitted with a normal distribution (black curve), Cross-sections of the black lines represent the mean value, and the range denotes the standard deviation. B) Number of SNAREpins formed in each phase was calculated separately and the ratios are plotted with the total number of SNAREpins formed. Green and orange dots represent Syp^30^VAMP2^60^Syt1^22^ and VAMP2^60^Syt1^22^ respectively. Syp-containing vesicles showed closer to 1:1 SNAREpin formation whereas VAMP2^60^Syt1^22^ vesicles were produced more randomly. C) Rate of SNAREpin assembly (number of SNAREpin formed per sec) in each phase is plotted with Poisson’s distribution fittings. Intersections of the black lines mark the mean values and ranges denote the standard errors whereas black squares represent the mean for the Poisson’s distribution. D) Table for detailed analyses of the number of SNAREpin formed and rate of assembly in each phase. n>45, n>3. * and ** p>0.01 and p>0.001 respectively.

**Figure S10.**
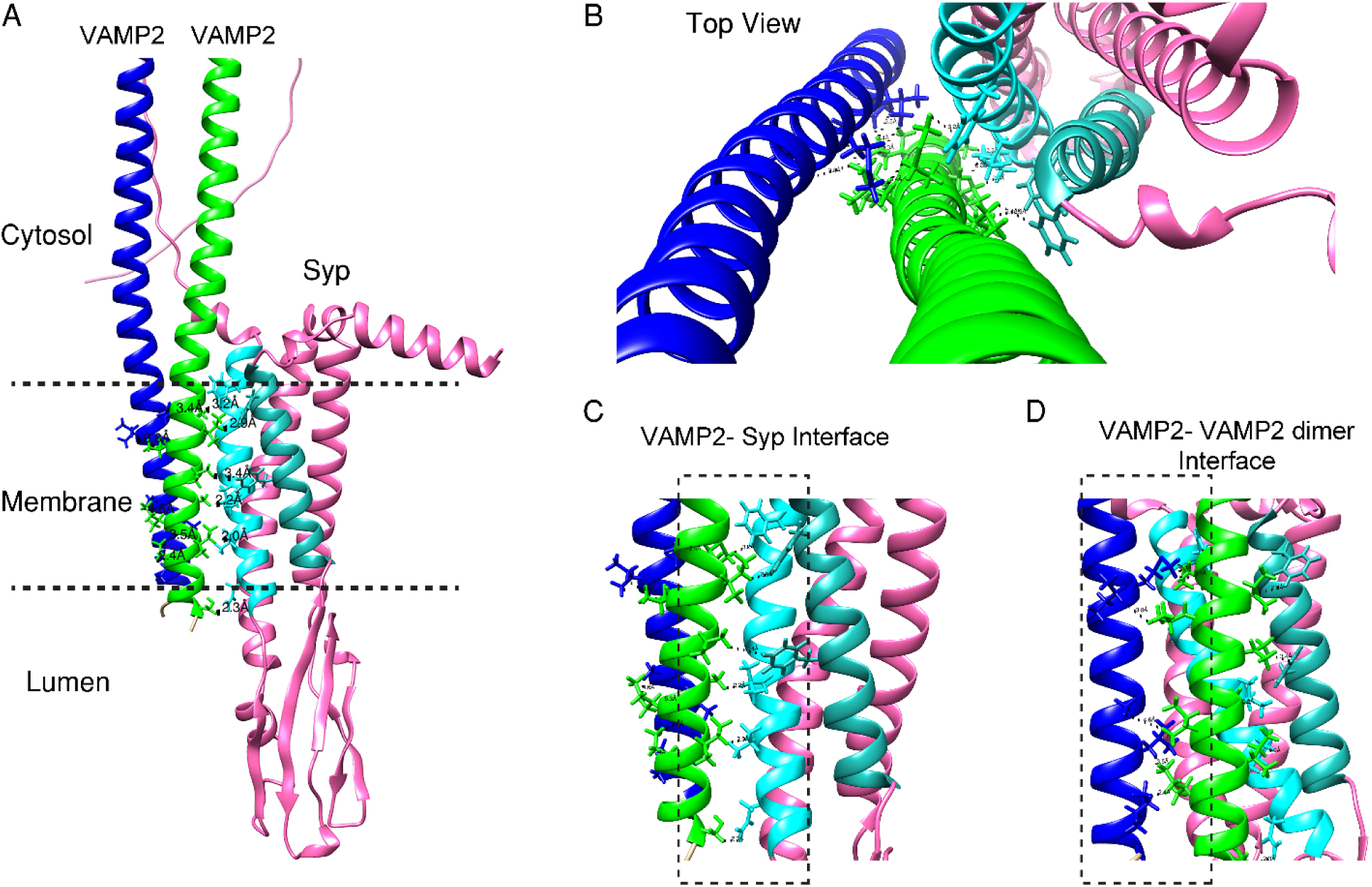
Prediction of VAMP2 dimer-Syp interaction by AlphaFold Colab2.3. A) Two VAMP2 (human) and one Syp (human) sequences were input and run for 20 recycles in AlphaFold Colab 2.3 (1). The output predicted two transmembrane domains of VAMP2 molecules can interact with the transmembrane domain of Syp with 70-90% confidence. Interestingly, one VAMP2 (green) is strongly predicted to interact with Syp 4^th^ (cyan) and 1^st^ (cornflower blue). The other VAMP2 (blue) has little interaction with Syp. Luminal and cytoplasmic domains are on opposite sides. B) Top view of the molecule where one VAMP2 (green) is interacting with both Syp and Vamp2. One interface of this molecule interacts with Syp. C) The Syp-VAMP2 interface could involve the following residues: 28Phe (Syp)-Ile97 (VAMP2) (distance 2.9 Å), 36Phe-101Val (3.4 Å), 210Leu-104Ala (2.2 Å), 206Val-108Ile (2.0 Å), 202Asn-115Ser (2.3 Å) and 221Phe-96 Met (3.2 Å). D) For the VAMP2-VAMP2 dimer interface: (blue-green VAMP2) 107Leu-110Ile (3.5 Å), 111Ile-110Ile (2.4 Å), 97Ile-99Leu (3.8 Å) and 104Ala-106Ile (4.6 Å). H-atoms were considered in distance calculations. Interacting residues are shown in sticks and black dotted lines. Chimera 1.16v (2, 3) was used to create these models and calculate the distances.

**Figure S11.**
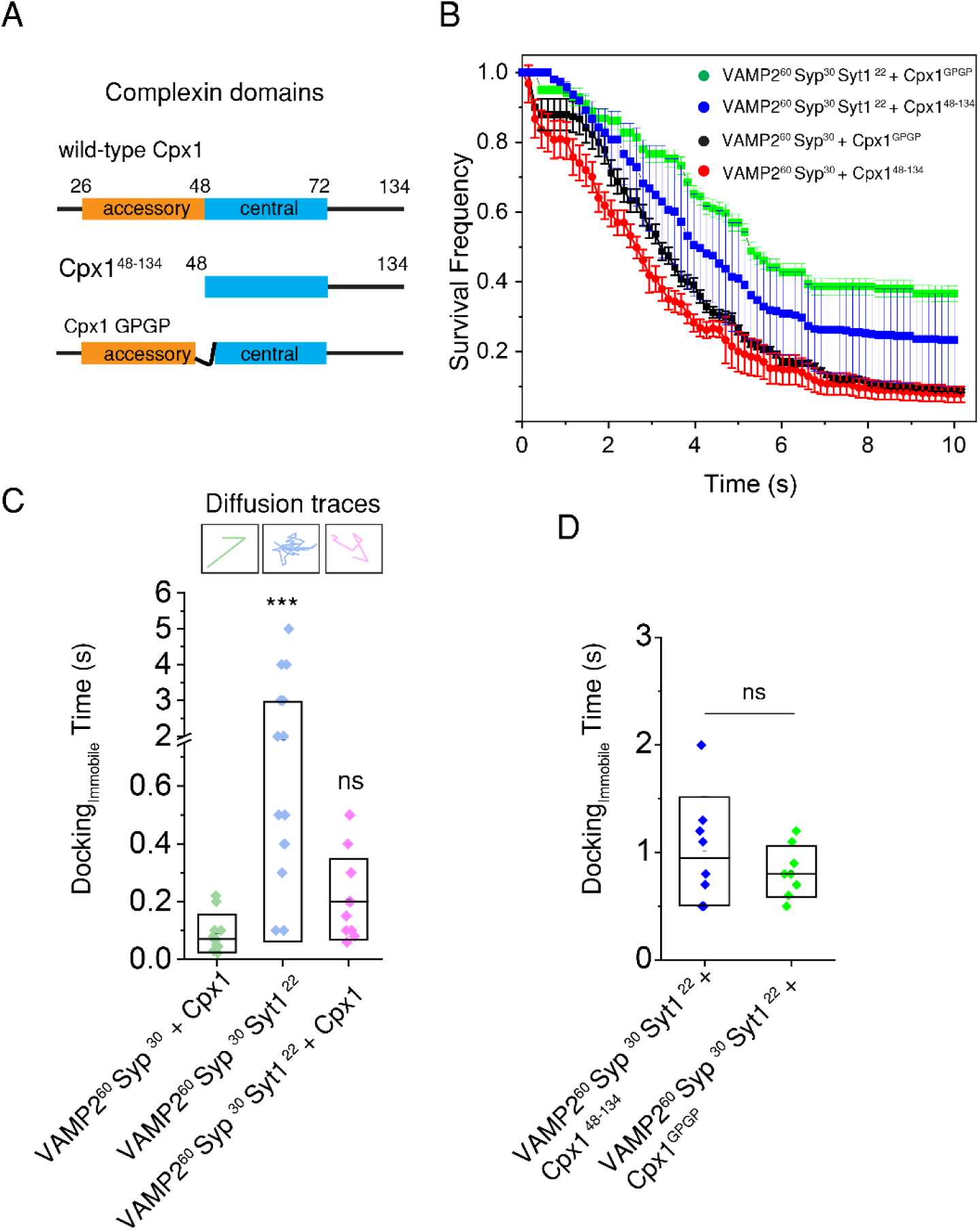
Complexin accessory helix and continuous helix are required for vesicle clamping. A) Schematic representation of Complexin domains. Residues 26-48 is the ‘accessory’ helix whereas 48-72 constitutes the ‘central’ helix. B) Survival analyses with Complexin mutants. Vesicles containing 60-copies of VAMP2 and 30-copies of Synaptophysin do not produce an efficient clamp in the presence of the accessory helix-deleted Cpx1^48-134^ (red) and helix-broken Cpx1^GPGP^ (black), at a Complexin concentration of 2 μM. An addition of 22-copies of Syt1 to these vesicles produce ~35% (with 2 μM Cpx1 ^48-134^, blue) and ~25% (with 2 μM Cpx1^GPGP^, green) clamping. These results suggest that the accessory helix and the central helix continuity are very crucial for efficient vesicle clamping. C) Docking time for immobile fractions is calculated where t=0 when the vesicle first appears in the movie series and t= docking time (immobile) when it first stably immobilizes on the bilayer. VAMP2^60^Syp^30^ vesicles in the presence of 2 µM Cpx1 (green) clamp within 90 ± 50 ms (mean ± SD). Interestingly, VAMP2^60^Syp^30^Syt1^22^ (blue) vesicles diffuse longer (1.5 ± 1.4 s) on the membrane before spontaneous fusion or stable clamping. However, the addition of Cpx1 to these vesicles (purple) accelerates docking time (0.2 ± 0.1 s). On top, individual diffusion traces are presented. Green, blue and purple colors represent the trace for VAMP2^60^Syp^30^ with Complexin, VAMP2^60^Syp^30^Syt1^22^ and VAMP2^60^Syp^30^Syt1^22^ with Complexin, respectively. Synaptophysin and Complexin act concomitantly to arrest these vesicles readily whereas Syt1 produces a little delay. For these experiments, the frame rate for data acquisition is 22 ms. * and *** represent P< 0.01 and 0.001, respectively. Each data point represents an average value from each independent experiment containing >25 vesicles. For each case, >100 vesicles are considered for final statistics. D) For both cases (with deletion of the accessory helix and the helix breaker mutants) vesicles are diffusing on the bilayer for ~1 s before becoming immobile. The frame rate for these experiments is ~150 ms. Each data point represents an average value for each independent experiment with ~15 vesicles, with total vesicles for each case >50. ns denotes not significant.

